# Network-based modeling of herb combinations in Traditional Chinese Medicine

**DOI:** 10.1101/2021.01.22.427821

**Authors:** Yinyin Wang, Hongbin Yang, Linxiao Chen, Mohieddin Jafari, Jing Tang

## Abstract

Traditional Chinese Medicine (TCM) has been practiced for thousands of years for treating human diseases. In comparison to modern medicine, one of the advantages of TCM is the principle of herb compatibility, known as TCM formulae. A TCM formula usually consists of multiple herbs to achieve the maximum treatment effects, where their interactions are believed to elicit the therapeutic effects. Despite being a fundamental component of TCM, the rationale of combining specific herb combinations remains unclear. In this study, we proposed a network-based method to quantify the interactions in herb pairs. We constructed a protein-protein interaction network for a given herb pair by retrieving the associated ingredients and protein targets, and determined multiple network-based distances including the closest, shortest, center, kernel, and separation, both at the ingredient and at the target levels. We found that the frequently used herb pairs tend to have shorter distances compared to random herb pairs, suggesting that a therapeutic herb pair is more likely to affect neighboring proteins in the human interactome. Furthermore, we found that the center distance determined at the ingredient level improves the discrimination of top-frequent herb pairs from random herb pairs, suggesting the rationale of considering the topologically important ingredients for inferring the mechanisms of action of TCM. Taken together, we have provided a network pharmacology framework to quantify the degree of herb interactions, which shall help explore the space of herb combinations more effectively to identify the synergistic compound interactions based on network topology.

## 1 Introduction

The pathogenesis and progression of many complex diseases are complicated such that the therapeutic effect of a single drug may be modest and further hampered by various side effects or drug resistance mechanisms^1^. Meanwhile, the pharmaceutical industry has begun to face the challenge of ‘more investments, fewer drugs’ in drug discovery. To reach the goal of better treatment efficacies and fewer side effects, there has been an increasing interest to investigate the synergistic effects of drug combinations^2^.

Although high-throughput phenotypic assays have been developed to screen potential drug combinations, an exhaustive search for the top hits from the huge combinatorial space arising from numerous agents remains a daunting task^3^. In contrast, computational approaches that leverage the rapid accumulation of pharmacological data may provide a cost-effective alternative to enable more systematic analyses of drug combinations. In particular, the advent of ‘omics’ technologies allow us to measure the drug perturbations in biological pathways and molecular interactions, resulting in an emerging systems-level approach called network pharmacology^4^. Instead of looking for one drug which acts solely on an individual target, multi-target drugs or drug combinations are more promising to achieve sustainable clinical response as many complex diseases have been shown to include multiple disease-causing genes^5-8^. In replacing the concept of ‘magic bullet’, this so-called network pharmacology paradigm requires accurate computational models that in many cases, can be used to predict an effective drug combination in order to perturb robustly disease phenotypes via targeting multiple pathways^9^. Ideally, such a drug combination should work synergistically to achieve stronger therapeutic effects with reduced doses of individual agents, so that the side effects may be minimized^9-11^.

To understand drug combinations better, we may look into an empirical paradigm of multi-component therapeutics known as Traditional Chinese medicine (TCM) to search for insights^12, 13^. Having been developed for over 3,000 years, TCM is characterized by the use of herbal formulae that usually consists of two or more medicinal herbs, which are capable of systematically preventing and treating various diseases via potentially synergistic herb interactions^14, 15^. Herb pairs involve a unique combination of two specific herbs, which form the most fundamental component of a multi-herb therapy^16^. By adding more herbs, a formula may be used to treat different diseases with greater flexibility^17-19^. For instance, *Coptis chinensis* (Huang Lian, used part: rhizome) and *Evodia rutaecarpa* (Wu Zhu Yu, used part: fruit) have been used together widely as formula ZuojinWan in clinical prescriptions for treating gastric diseases as a basic herb pair ^20^. Depending on the additional herbs that are mixed with *Coptis chinensis* and *Evodia rutaecarp*, they have been used for many disease indications, including the inhibition of inflammation^*21*^, as well as treating hypertension^22^ and obesity^23^.

Considering the important role of herb pairs in the development of TCM, it might be of great significance to investigate the rationale of why certain herb pairs are commonly used for treating a particular disease^24, 25^. However, there exists very limited understanding at the molecular level on how the herb pairs work synergistically to achieve stronger therapeutic effect^12, 26, 27^. One of the major bottlenecks is that herb combination is inherently more complex as herbs usually consist of multiple ingredients. Recent studies suggested that synergistic effects in herb combinations mainly rely on the interactions of their ingredients, leading to boosted treatment effects compared to single herbs^28^. One example is the cardio-protective effects by the combination of Paeonol (isolated from the root cortex of the *Paeonia moutan [Syn Paeonia suffruticosa]*) and Danshensu (isolated from the root of the Chinese herb *Salvia miltiorrhiza*)^29^. Another example is the comination of *icariin* from *aerial parts* of herb *Epimedium brevicornum* (Yin Yang Huo*), berberin* from the bark of *Phellodendron amurense* (Huang Bai), and *curculigoside* from rhizome of *Curculigo orchioides* (Xian Mao) in the Er-Xian decoction, which can produce synergistic effects on Osteoclastic bone resorption^30^. Furthermore, ingredients within an herb might also interact synergistically to induce pharmacological effects. One example is the interaction of *ginsenoside Rb1*, g*insenoside Rg1* and *ginsenoside 20(S)-protopanaxatriol* found in the root of *Panax ginseng*, which can produce synergistic effects on their antioxidant activity^31^. These individual studies on specific herbs form the basis for developing a more systematic method to model the interactions among TCM herbs at the molecular level, which may hold the key to rationalize the herb combinations for future drug discovery.

Recently, network pharmacology approaches have been introduced for the study of drug interactions for a variety of diseases^32, 33^. For example, Huang *et al*. proposed a novel tool called DrugComboRanker based on drug functional network to prioritize potential synergistic drug combinations and further validated their mechanisms of action in lung adenocarcinoma and endocrine receptor positive breast cancer^34^. Cheng *et al*. proposed a network-based methodology to characterize the distance between two drugs according to their target distributions in a protein-protein network^11^. They demonstrated that clinically approved drug combinations tend to have lower distance compared to random drug pairs, and for a drug pair working synergistically for a given disease, both of them need to hit the disease module but via non-overlapping network neighborhood. Furthermore, a modularity analysis of multipartite networks has suggested that network modeling might be a promising method for understanding the mechanisms of actions of traditional medicine^35^. With the great success in understanding the interaction between chemicals and diseases, network-based models warrant further studies to make sense of the rationale of TCM herb interactions.

In this study, we hypothesized that network pharmacology models on the underlying drug-target interactions behind the herb combination may provide novel insights into herb pair’s mechanisms of action, which are critical for the phenotypic-based drug discovery from TCM^36-38^. We investigated the frequencies of herb pairs that appear in the common TCM herb formulas. We developed a network-based model to characterize the distance of herbs within an herb pair in a protein-protein interaction network. The model considered the interactions of herbs at the herb, ingredient and target levels, and utilized five distance metrics including the closest, shortest, separate, kernel and center methods. In addition, Area under curve (AUC) of precision and recall (PR) as well as receiver operating character characteristics (ROC) were used to determine the best distance metric for discriminating the most frequent herb pairs against non-existing herb pairs. Finally, we found that a commonly used herb pair tends to have smaller network distance compared to non-existing herb pairs, suggesting that herb combinations tend to achieve stronger protein-protein interactions. In addition, we found that the center ingredients of herbs tend to play important roles. In a case study of an herb pair including *Astragalus membranaceus* and *Glycyrrhiza uralensisn*, we further showed that their network-based distance is significantly smaller than random and then center ingredients of the herb pair. Taken together, the network modeling approach provides a more systematic framework to characterize herb interactions at the molecular level that may lead to the rationalization and modernization of TCM herb combinations ultimately^27^.

## 2 Methods

### 2.1 Collection of herb pairs

We searched for existing herbal formulae from TCMID, a manually-curated TCM database^39^. TCMID is by far one of the most comprehensive TCM databases. More importantly, compared to other databases, TCMID supports data download service, which facilitates the effective integration of TCM data and PPI data in our study. Therefore, for allowing a more systematic analysis of the TCM herbs, we decided to use the data from TCMID. There are 8159 herbs and more than 25,210 herb ingredients in the TCMID database in total. However, after filtering out herbs and ingredients that are lack of target information, 349,197 herb pairs were collected from 46,929 herbal formulae, including 4415 herbs, 4330 ingredients, 3171 targets, 17,753 herb-ingredient pairs as well as 25,050 ingredient-target pairs. As the same herb pair may appear in multiple herbal formulae, we considered the top 200 most frequent herb pairs with target information for both herbs (frequencies between 358 and 3846) as a positive set (**Supplementary Figure 1**). In contrast, we determined 10,000 randomly generated herb pairs, out of which we considered 9459 herb pairs that were not observed in the actual herbal formulae as a negative control data set. Therefore, the positive set represents the common herb pairs while the negative set represents the herb pairs that are not used in any of the herbal formulae. To obtain an independent validation set, we also collected 268 herb pairs that have been considered as basic components of herbal formulae according to traditional medicine literature^14, 16, 40^.

### 2.2 Extraction of interactions between herbs, ingredients and targets

We collected the herb-ingredient information from the TCMID. Herbs that lack ingredient information were not considered. Similarly, ingredient compounds without structural information were discarded, as they could not be modelled in the PPI network analysis. For the remaining ingredient compounds, their targets were extracted from the STITCH database^41^. Target-target interactions were extracted from a manually-curated human interactome including 243,603 PPIs and 16,677 proteins^11^, which are assembled from commonly-used databases including IntAct^42^, InnateDB^43^, PINA^44^, HPRD^45^, BioGRID^46^, HI-II-14_Net^47, 48^, PhosphositePlus^49^, KinomeNetworkX^50^, INstruct^51^, SignaLink2.0^52^ and MINT^53^. These databases cover a wide range of protein-protein interaction data derived from experimental and computational approaches. All the interactions were denoted as undirected edges in the network.

### 2.3 Network proximity models of herb pairs

The herb-herb distance can be determined by considering the ingredients as the nodes, where for a pair of ingredients their distance can be further determined from their target profiles in the PPI network. Denote that *I*(*A*) = (*a*_1_, *a*_2_, …) is the ingredient set for a herb *A*, where for an ingredient *a* the set of targets is *T*(*a*) = (*t*_1_, *t*_2_, …). For another herb *B*, its ingredient set and target sets are defined similarly. We applied five measures introduced by Cheng *et al*.^11^ to determine the network distance between two herbs, including closest, separation, shortest, kernel and center.

The closest distance is defined as:

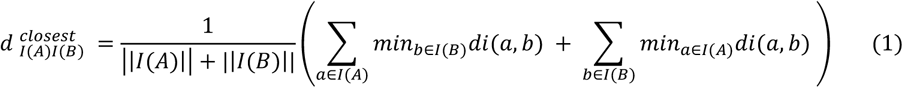

where *di* (*a, b*) is the distance between two ingredient nodes in herb *A* and herb *B*, and ||*I(A)*|| and ||*I(B)*|| are the numbers of ingredients for herb *A* and *B*, separately. For each ingredient in herb A, we considered its distance with all the ingredient nodes in herb B, and determined the minimal distance as its closest distance. As shown in equation (1), we determined the mean closest distance for all the ingredients in *A* and *B*, and used it as the closest distance 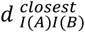 between the two herbs.

The separation distance is defined as the closest distance between *A* and *B*, subtracted by the average closest distances within *A* and *B*:

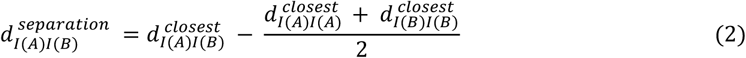

The shortest distance sums up all the distances between nodes in *A* and *B*, and then normalized by the product of their sizes:

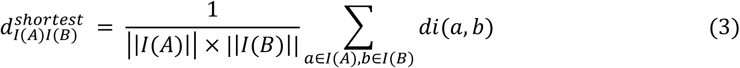

The kernel distance is defined as the average of exponent-based pairwise distance, normalized by their relative network sizes:

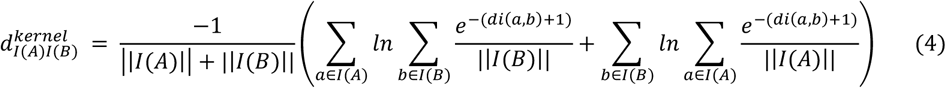

The center distance identifies the centers of A and B as the nodes with minimal sum of distances, and then determines the distance between the two centers:

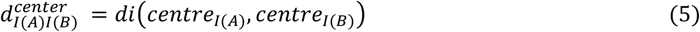

where

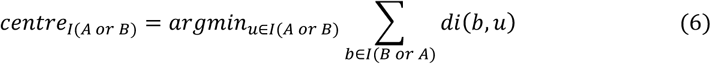

The equations (1-6) involve the calculation of distances for two ingredients (*a, b*), for which we again have five options based on their target profiles *T(a)* and *T(b)* including:

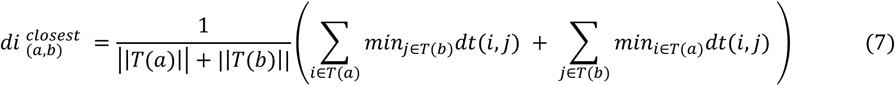

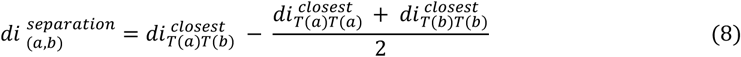

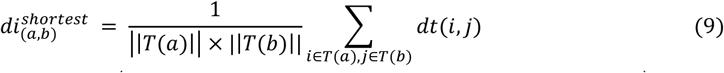

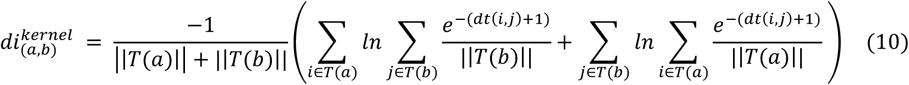

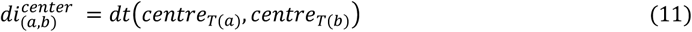

As we considered five distance methods that can be applied at both the target and the ingredient levels, the network proximity can be defined by an exhaustive combination of them, resulting in 25 distance models in total. For example, a model can be constructed using *closest (ingredient) - closest (target)* distance, defined as the closest distance for two herbs at the ingredient level:

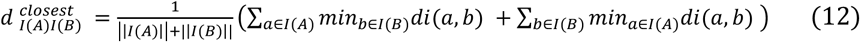

where *di* (*a, b*) for ingredient *a* and ingredient *b* is:

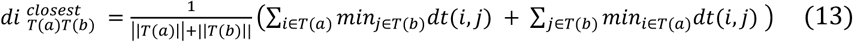

where *dt* (*i, j*) is the shortest path length between the two targets in the PPI network^54^.

### 2.4 Discrimination performance of the proximity distances

We utilized the area under the receiver operating character characteristic (ROC) curve (AUC) to evaluate discriminative ability of the network proximity models for separating the top frequent herb pairs and non-observed random herb pairs. True positive rate and false positive rate were determined at different thresholds of network proximity value. To obtain a balanced data set with an equal number of positive and negative cases, we randomly selected two herbs as non-observed herb pairs from the 4415 herbs for 200 times, resulting in a set of 200 negative herb pairs for comparison. To determine the average AUC scores, we repeated the procedure 50 times. For the 268 literature-mined herb pairs (described in section 2.1 as an independent validation set), we also repeatedly generated 268 random pairs as negative control.

### 2.5 A case study on modeling the combination of *Astragalus membranaceus* and *Glycyrrhiza uralensis*

It is reported that the herb pair Huang Qi (the root of *Astragalus membranaceus*) and Gan Cao (the root and rhizome of *Glycyrrhiza uralensis*) can be used for liver fibrosis and cirrhosis treatment, while neither *Astragalus membranaceus* nor *Glycyrrhiza uralensis* shows therapeutic effects when used alone^55, 56^. Therefore, it is important to identify the synergistic interactions of the ingredients underlying the herb pair for treating liver diseases. To explore the mechanisms of the herb pair, we constructed the herb-herb network based on their ingredients and targets. We first evaluated whether the distance between *Astragalus membranaceus* and *Glycyrrhiza uralensis* is different from the expectation of a random herb pair. Furthermore, we identified the center ingredients that are more likely to explain the synergy of the two herbs. Finally, we performed pathway analysis using enrichr^57^ based on the target genes of the center ingredients.

## 3 Result

### 3.1 Frequency of single herbs and herb pairs

There are 8159 herbs and more than 25210 herb ingredients in the TCMID database in total. However, after filtering out herbs and ingredients that lack target information, 349,197 herb pairs were collected from 46,929 herbal formulae, including 4415 herbs, 4330 ingredients, 3171 targets, 17,753 herb-ingredient pairs as well as 25,050 ingredient-target pairs. Most of the herb formulae (97.9 %) contain less than 20 herbs, with an average of 4.93 (**Supplementary Figure 1**). The herbs with top-ten highest frequencies are Gan Cao (root and rhizome of *Glycyrrhiza uralensis*, 12518), Dang Gui (root of *Angelica sinensis*, 7417), Ren Shen (root of *Panax Ginseng*, 7390), Bai Zhu (5259, root of *Atractylodes macrocephala [Syn. Atractylis macrocephala]*), Huang Qin (4163, root of *Scutellaria baicalensis*), Fang Feng (4074, root of *Saposhnikovia divaricata [Syn. Ledebouriella seseloides])*, Chuan Xiong (4007, rhizome of *Ligusticum chuanxiong [Syn. Ligusticum wallichii]*), Fu Ling (3666, sclerotium of *Poria cocos*), Chen Pi (3650, from the dried peel of *Pericarpium Citri Reticulatae*) (**Supplementary Table 1**). *Glycyrrhiza uralensis* is extensively used as a major component in the 12,518 prescriptions, supported by its various pharmacological activities including anti-inflammatory, anti-oxidative, antidiabetic, hepatoprotective and memory enhancing activities^58^. *Angelica sinensis* is widely applied for menstrual disorders by enhancing the blood circulation, and also has been reported to have multiple immunomodulation and anti-inflammation, as well as cardio-cerebrovascular effects^40^. *Panax Ginseng* is commonly used as a functional food with a long medical history, which has shown efficacy in multiple diseases, such as anti-cancer, neurodegenerative disorders, insulin resistance and hypertension. Another important effect of *Panax Ginseng* is maintaining homeostasis of the immune system^59-61^. All the top three most frequent herbs tend to activate the immune system, suggesting the importance of activating the immune system when prescribing TCM. This observation is consistent with the TCM theory, where these herbs are usually called tonifying (adjuvant) herbs that possess supplementing and strengthening the treatment effects in addition to the major herbs.

These high-frequent herbs also tend to show higher chances to be combined with the other herbs (**Figure 2**). For example, *Panax Ginseng* and *Glycyrrhiza uralensis* appear together in 3846 of 46,929 herbal formulae, followed by the pair of *Angelica sinensis* and *Glycyrrhiza uralensis* that are co-administered in 2907 herbal formulae. However, the majority of the 349,197 herb pairs (99.4%) occurred in less than 100 herbal formulae. Only 163 herb pairs of the remaining 1950 (0.6%) herb pairs showed a frequency higher than 500 (**Supplementary Figure 2**).

**Figure 1:**
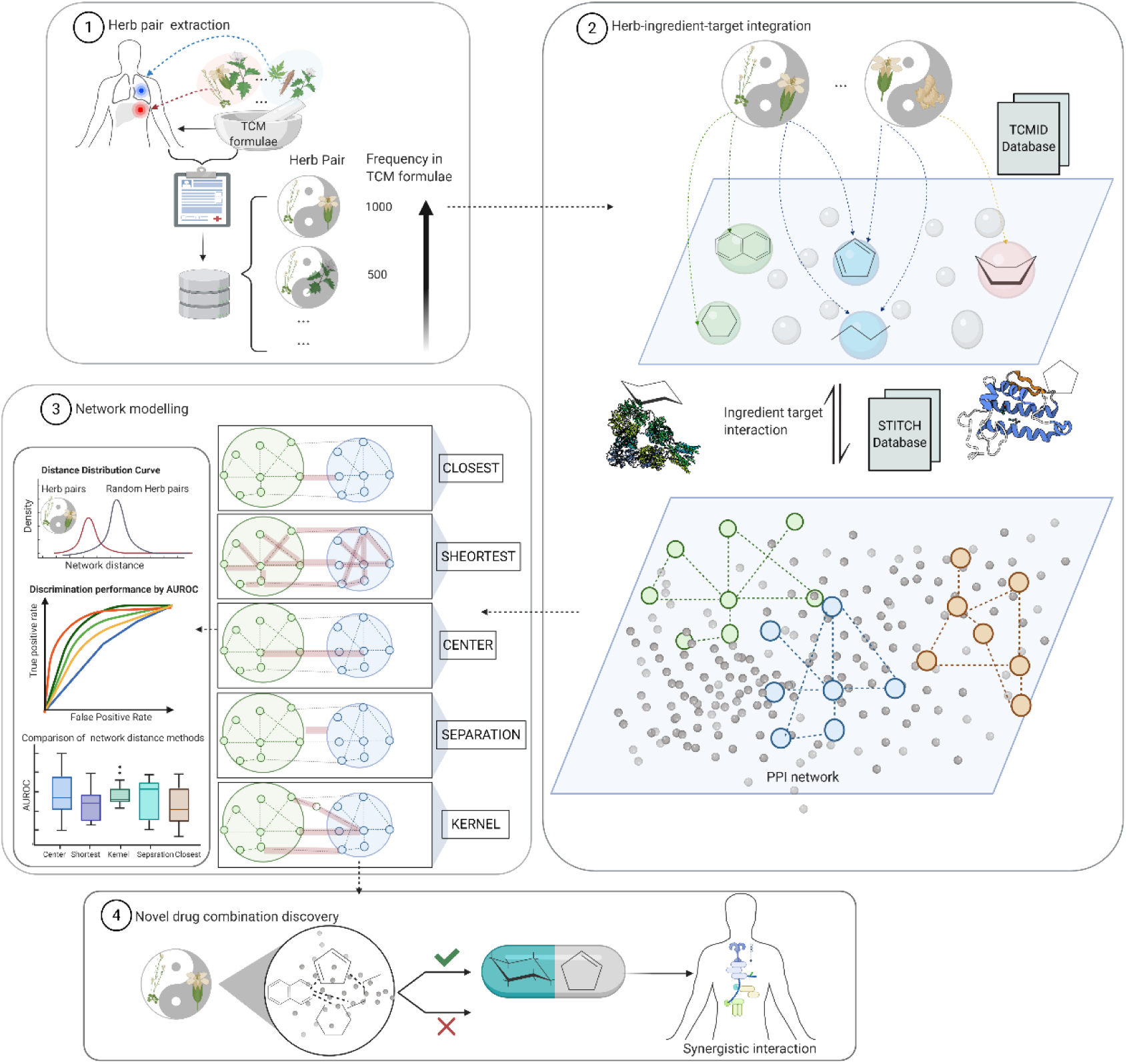
Workflow of the network construction for herb pairs. Top frequent herb pairs were determined from existing herbal formulas. For each herb pair, the network consists of three levels of interactions including herb-ingredient, ingredient-target and target-target interactions. The network proximity can be determined at either the ingredient level or the target level by multiple metrics including the closest, shortest, separate, kernel and center distances. We aimed to determine the network models that can separate the most frequent from the least frequent herb pairs.

**Figure 2:**
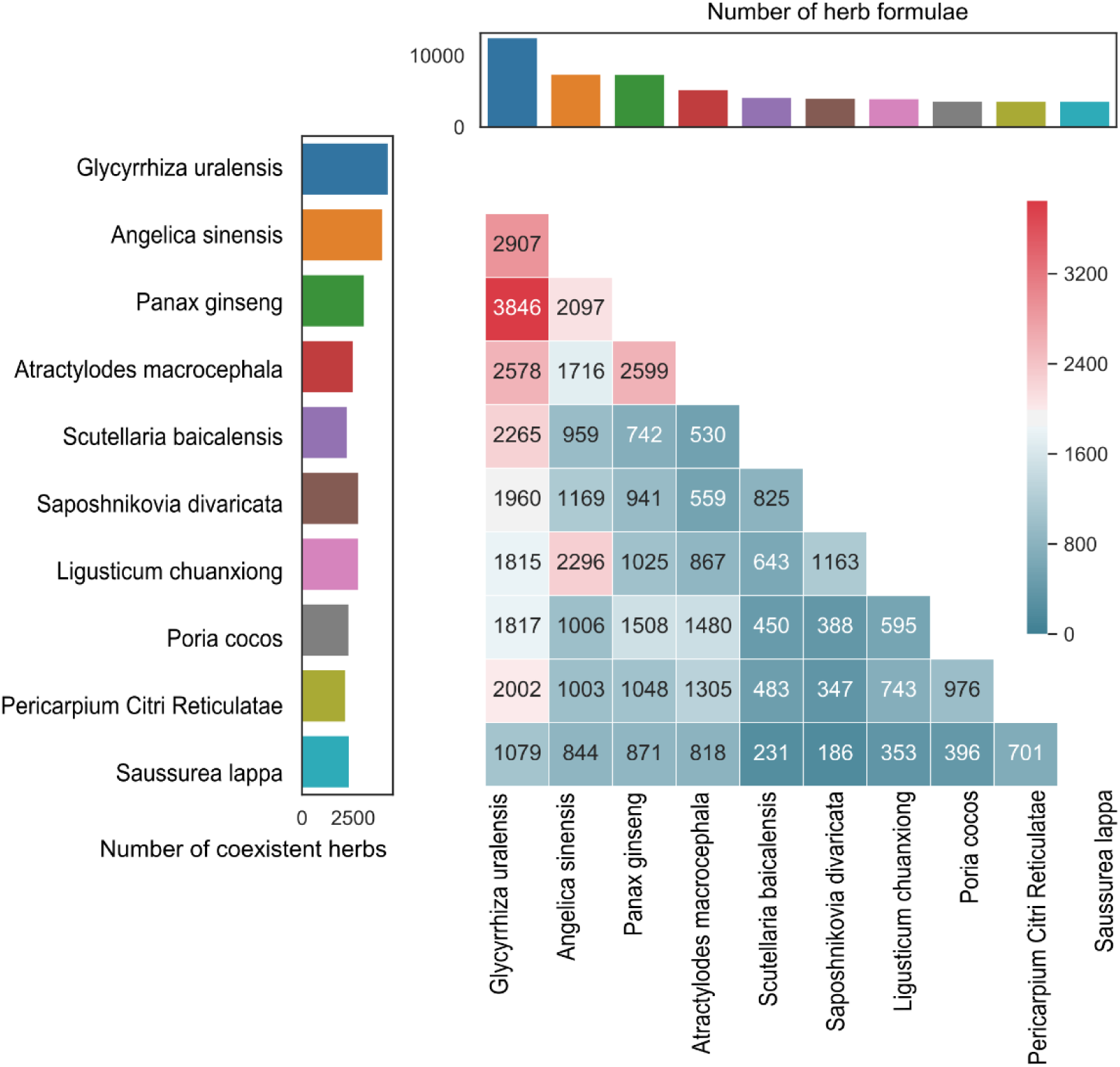
Patterns of pairing for the top ten most frequent herbs. The frequency of the herbs is shown in the top panel while the number of unique herbs that are co-administrated with them is shown in the left panel. The numbers inside the heat map show the frequencies of their pairwise combinations.

As shown in the **Supplementary Figure 2**, there is a sharp decrease of herb pair frequency after 200. Therefore, we considered those herb pairs with frequency larger than 200 and target information for both herbs to be the popular herb pairs. In the following analyses, we focused on these top herb pairs and searched for their target and ingredient information (**Supplementary Table 2**). These herb pairs involve 61 unique herbs, for which the average number of ingredients is 16.80. There is at least one common ingredient for 43% (86) of the top 200 herb pairs, while only 2.08% of randomly generated herb pairs share at least one ingredient (**Supplementary Figure 3**). Use of common ingredients tends to be a strategy of TCM prescription, as it was found that synergistic effects may be achieved by affecting the same pathways with common or similar compounds^62^. For example, Qiang Huo (the rhizome or root part of *Notopterygium incisum*) and Du Huo (the root part of *Angelica pubescens f. biserrata [Syn. Angelica pubescens])* share ten common ingredients (including *gamma-amin*.*yri*., *camphor, columbianetin, guaiol, guanidinium, isoimperatorin, isopimpinellin, nodakenin, scopoletin, and osthole*) and have appeared in 522 herbal formulas. At the same time, different ingredients in these herb pairs may play various roles, such as optimization of pharmacodynamics and/or pharmacokinetics to improve therapeutic efficacy and/or reduce toxicity and adverse reactions^17^, which can be explained by the “Jun-Chen-Zuo-Shi” theory in TCM system^63^. For example, the combination of *cacalol* from plant *Cacalia delphinifolia* and *paclitaxel* extracted from the yew trees can significantly suppress tumor growth and overcome chemo-resistance^64^.

### 3.2 Network distance for top-frequent herb pairs

We modelled the interactions for an herb pair at two levels including the ingredient and the target levels. For each level, we considered five distance methods including closest, separation, shortest, kernel and center. In the next step, we examined all combinations of distance metrics in both levels, resulting in 25 (5*5) distance models in total. We focused on the top 200 most frequent herb pairs and determined their network-based distances, as compared to randomly selected herb pairs. We found that the average network distance of the top herbs pairs is mostly less than the average distance of random herb pairs, with statistical significance in 16 of the 25 distance models (p-value <0.05) (**Figure 3, Table 1**). For example, the center-separation model showed the best performance to differentiate the top herb pairs from random pairs, with a difference of 0.489 (p-value = 9.91E-28, t-test). As the herb-herb network is constructed based on their interactions in ingredients and targets, a shorter distance therefore indicates that herb pairs tend to affect similar pathways in order to produce synergistic effects. We also examined the likelihood of a top-frequent herb pair sharing the same ingredients, which might explain why they have shorter distance. These shared ingredients may contribute partly to the closer distances of the herb pairs.

**Table 1:**
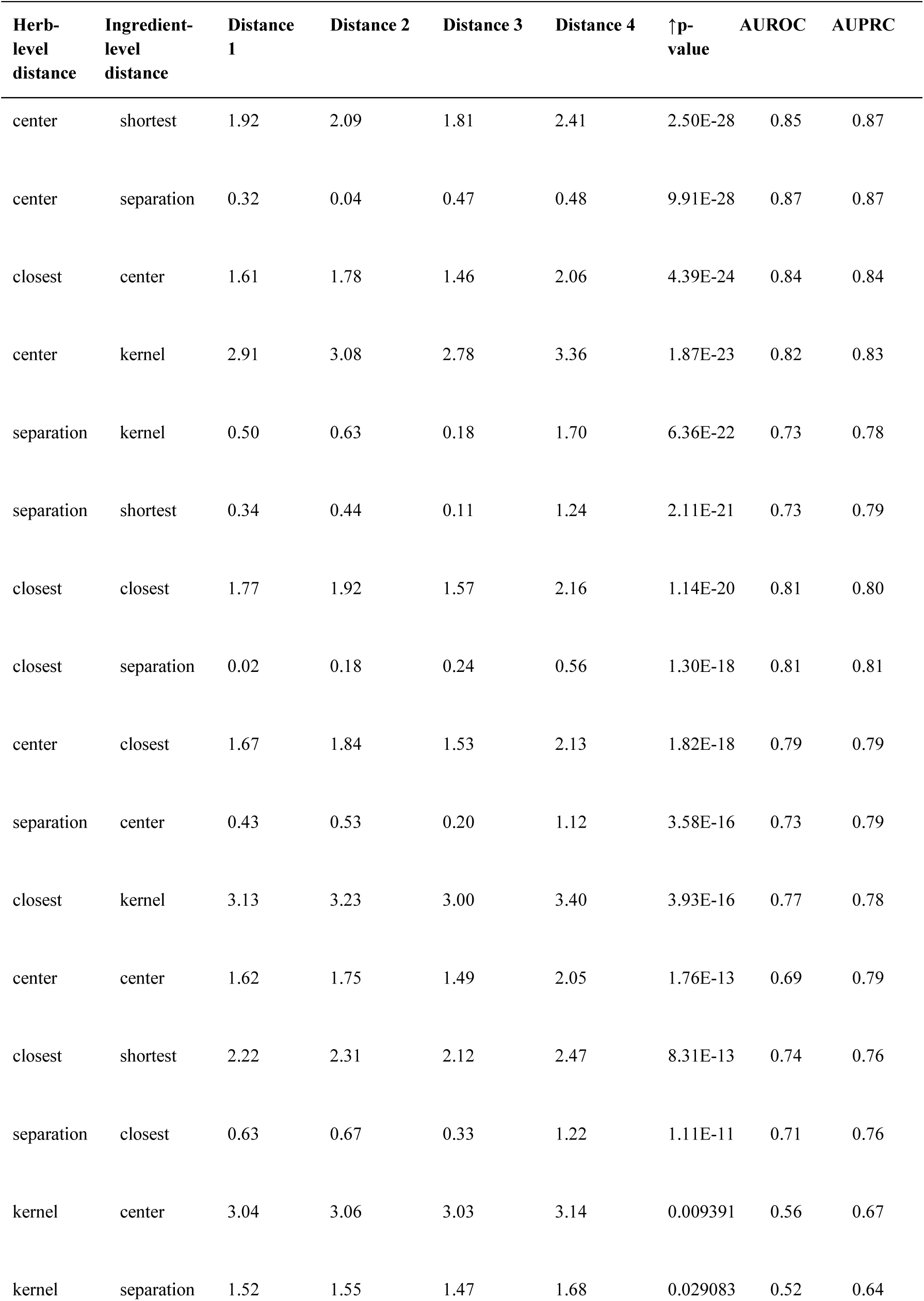
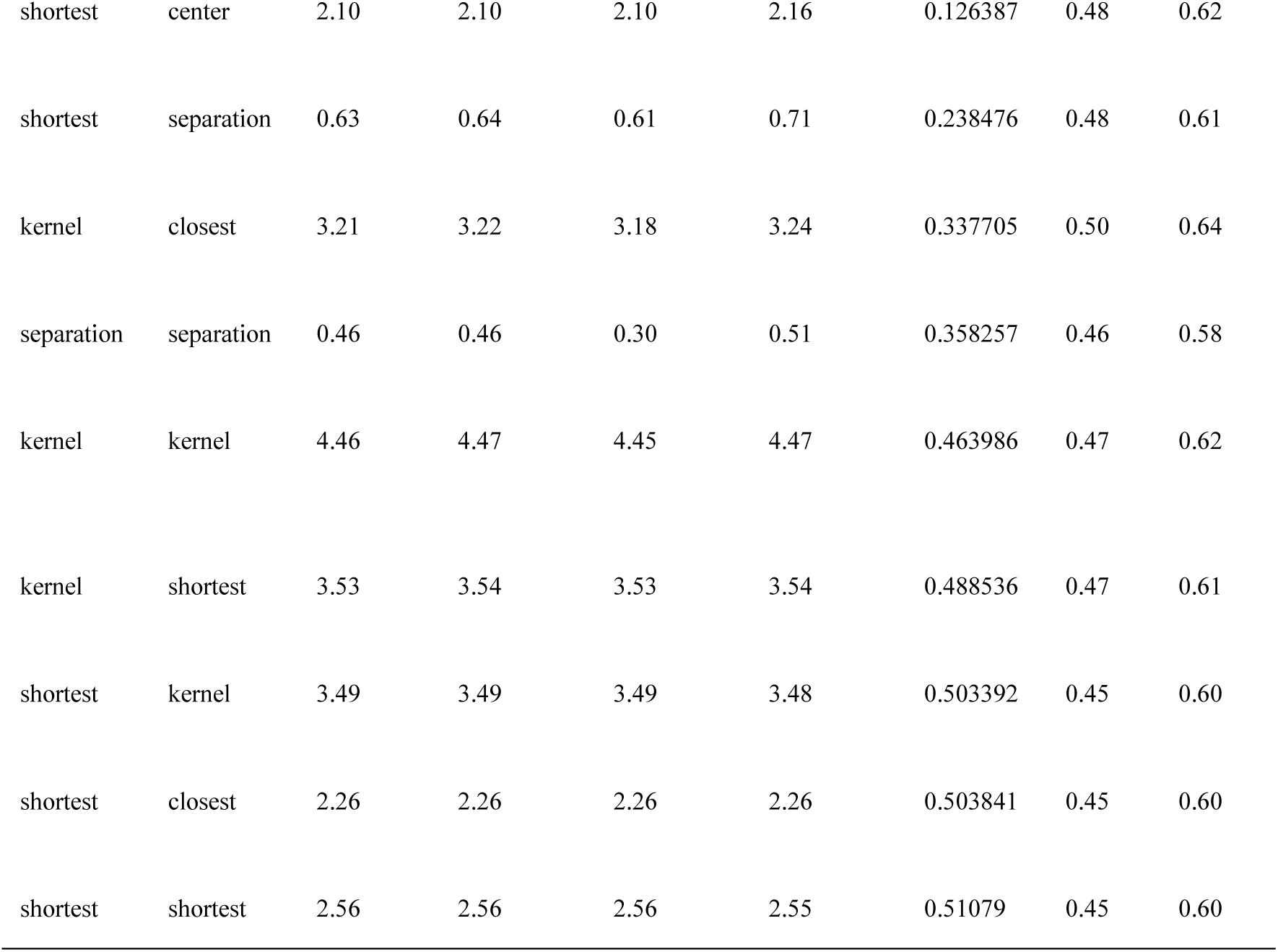
Comparing the network proximity models. The p-values are determined by the difference between the top 200 herb pairs and random herb pairs. Distances 1, 2, 3, 4 are the average distance for top 200 herb pairs, top 10000 herb pairs, top non-overlapping 114 herb pairs and random herb pairs, respectively.

| Herb-level distance | Ingredient-level distance | Distance 1 | Distance 2 | Distance 3 | Distance 4 | ↑p-value | AUROC | AUPRC |
| --- | --- | --- | --- | --- | --- | --- | --- | --- |
| center | shortest | 1.92 | 2.09 | 1.81 | 2.41 | 2.50E-28 | 0.85 | 0.87 |
| center | separation | 0.32 | 0.04 | 0.47 | 0.48 | 9.91E-28 | 0.87 | 0.87 |
| closest | center | 1.61 | 1.78 | 1.46 | 2.06 | 4.39E-24 | 0.84 | 0.84 |
| center | kernel | 2.91 | 3.08 | 2.78 | 3.36 | 1.87E-23 | 0.82 | 0.83 |
| separation | kernel | 0.50 | 0.63 | 0.18 | 1.70 | 6.36E-22 | 0.73 | 0.78 |
| separation | shortest | 0.34 | 0.44 | 0.11 | 1.24 | 2.11E-21 | 0.73 | 0.79 |
| closest | closest | 1.77 | 1.92 | 1.57 | 2.16 | 1.14E-20 | 0.81 | 0.80 |
| closest | separation | 0.02 | 0.18 | 0.24 | 0.56 | 1.30E-18 | 0.81 | 0.81 |
| center | closest | 1.67 | 1.84 | 1.53 | 2.13 | 1.82E-18 | 0.79 | 0.79 |
| separation | center | 0.43 | 0.53 | 0.20 | 1.12 | 3.58E-16 | 0.73 | 0.79 |
| closest | kernel | 3.13 | 3.23 | 3.00 | 3.40 | 3.93E-16 | 0.77 | 0.78 |
| center | center | 1.62 | 1.75 | 1.49 | 2.05 | 1.76E-13 | 0.69 | 0.79 |
| closest | shortest | 2.22 | 2.31 | 2.12 | 2.47 | 8.31E-13 | 0.74 | 0.76 |
| separation | closest | 0.63 | 0.67 | 0.33 | 1.22 | 1.11E-11 | 0.71 | 0.76 |
| kernel | center | 3.04 | 3.06 | 3.03 | 3.14 | 0.009391 | 0.56 | 0.67 |
| kernel | separation | 1.52 | 1.55 | 1.47 | 1.68 | 0.029083 | 0.52 | 0.64 |
| shortest | center | 2.10 | 2.10 | 2.10 | 2.16 | 0.126387 | 0.48 | 0.62 |
| shortest | separation | 0.63 | 0.64 | 0.61 | 0.71 | 0.238476 | 0.48 | 0.61 |
| kernel | closest | 3.21 | 3.22 | 3.18 | 3.24 | 0.337705 | 0.50 | 0.64 |
| separation | separation | 0.46 | 0.46 | 0.30 | 0.51 | 0.358257 | 0.46 | 0.58 |
| kernel | kernel | 4.46 | 4.47 | 4.45 | 4.47 | 0.463986 | 0.47 | 0.62 |
| kernel | shortest | 3.53 | 3.54 | 3.53 | 3.54 | 0.488536 | 0.47 | 0.61 |
| shortest | kernel | 3.49 | 3.49 | 3.49 | 3.48 | 0.503392 | 0.45 | 0.60 |
| shortest | closest | 2.26 | 2.26 | 2.26 | 2.26 | 0.503841 | 0.45 | 0.60 |
| shortest | shortest | 2.56 | 2.56 | 2.56 | 2.55 | 0.51079 | 0.45 | 0.60 |

**Figure 3.**
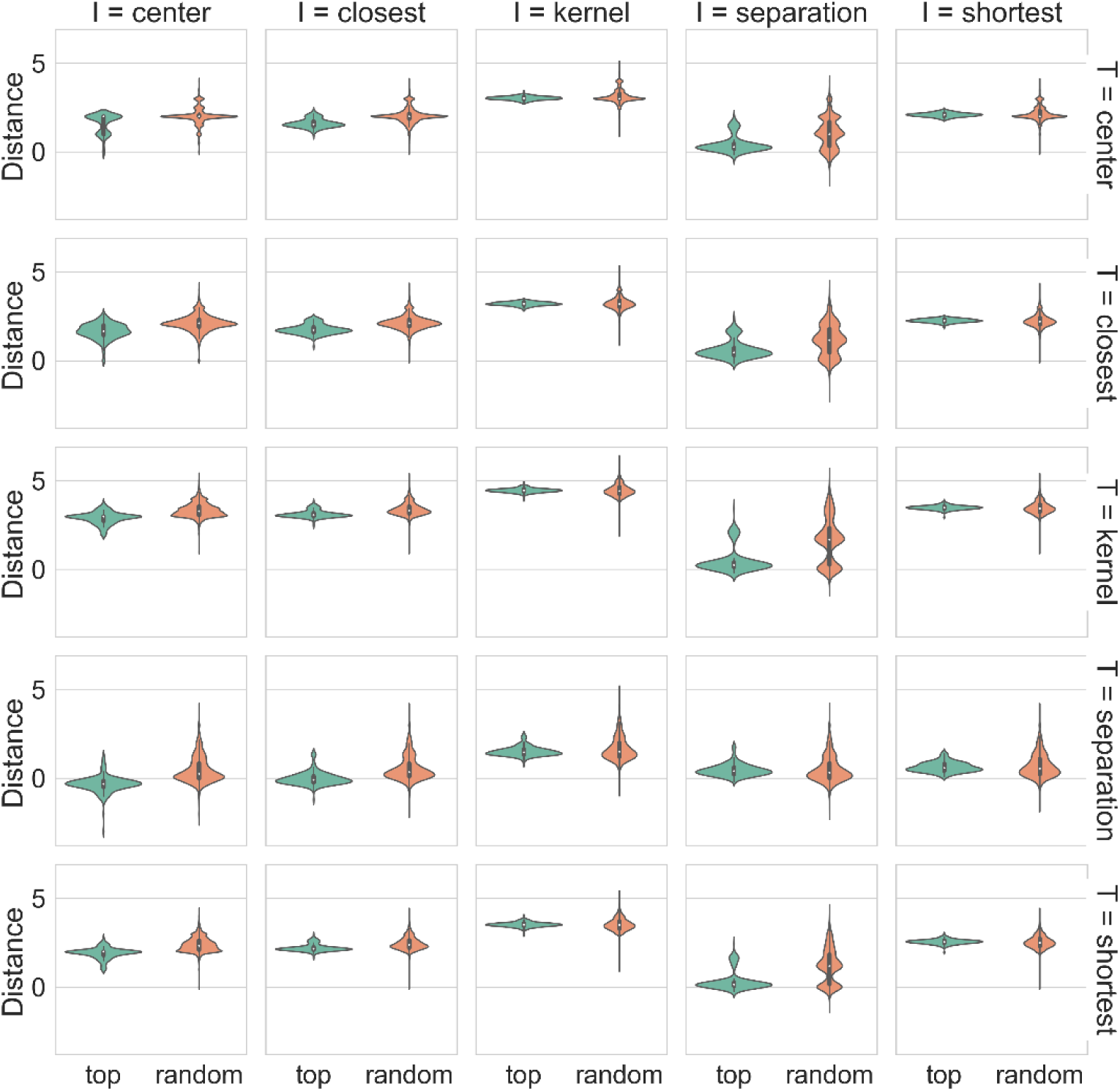
Network distances for the top herb pairs when comparing with random herb pairs. ‘I’ stands for the ingredient-level distance methods and ‘T’ stands for the target-level distance methods.

We found that 114 out of the 200 herb pairs did not share any common ingredients, while a few herb pairs (n = 15) shared more than three ingredients **(Supplementary Figure 3**). However, when we considered the 114 herb pairs that did not share any common ingredients, we still found that their distances are significantly lower than that for random herb pairs **(Supplementary Figure 4)**. This result suggested that in addition to the common ingredients, target interactions from different ingredients within an herb pair remain a major mechanism of action to affect functionally related disease pathways.

### 3.3 Discrimination performance of the distance metrics

To evaluate the discrimination power of the network models, we determined the Receiver Operating Characteristic (ROC) curve and Precision-Recall (PR) curve using the top frequent herb pairs as positive cases and random herb pairs as negative cases. In general, we found that the average AUROC (Area Under the ROC curve) and AUPRC (Area Under the PR curve) for the 25 distance metrics reach 0.65 and 0.72, respectively, suggesting the general validity of using the network-based distance metrics to characterize the herb-pair interactions (**Table 1**). We found that the top performance was achieved by two models that utilize the center distance at the ingredient level, including the center (ingredient) - separation (target) model and the center (ingredient) - shortest (target) model. The ROC curves for these two models were shown in **Figure 4**, confirming the superior discrimination performance.

**Figure 4.**
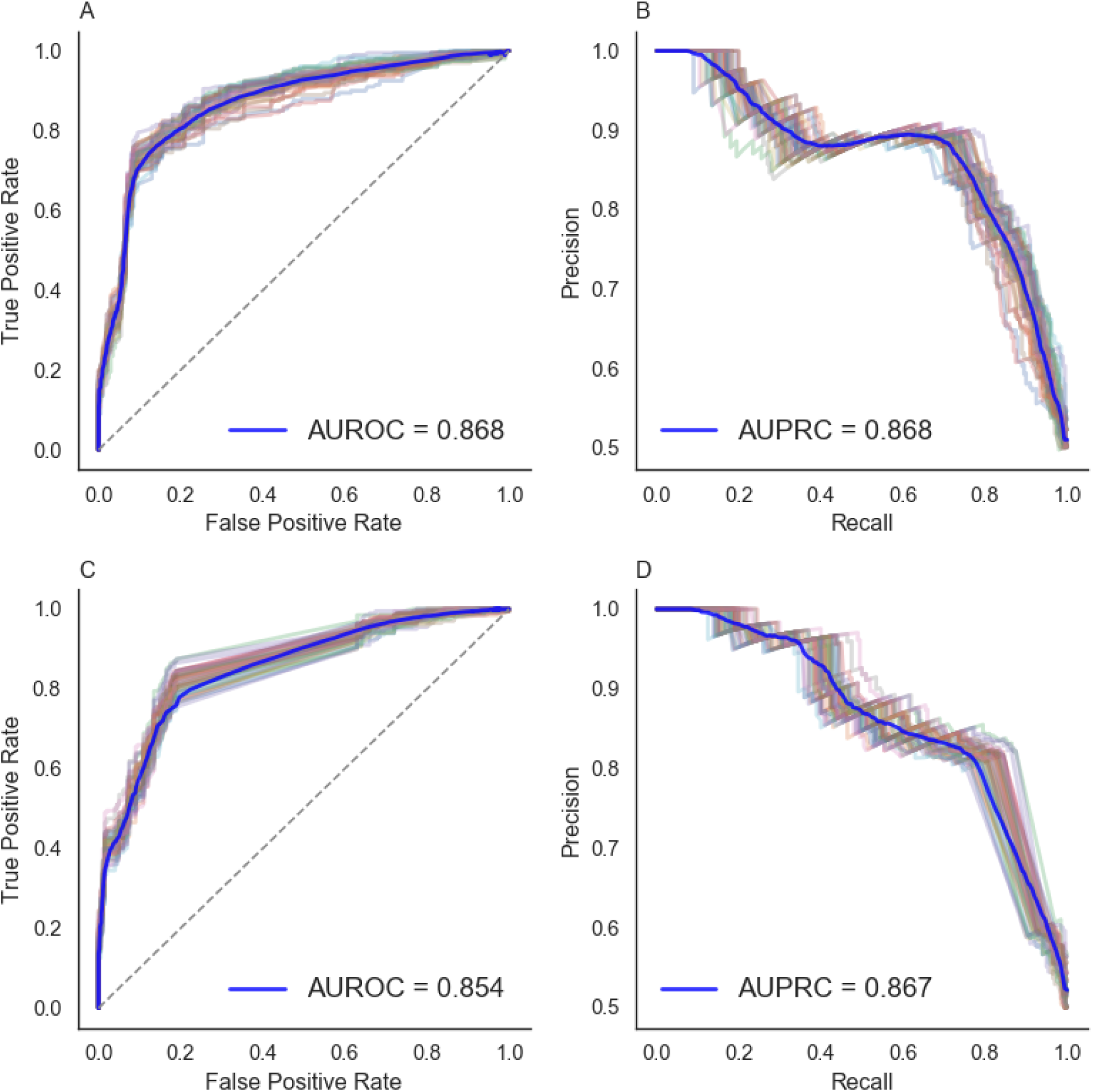
Receiver operating characteristic curves and precision-recall curves of the (A-B) center-separation model and (C-D) the center-shortest model. Each curve is a result of one permutation while the blue curve is the average value of all the permutations.

Interestingly, we found that the five models utilizing the center distance at the ingredient level (i.e. center (ingredient) - center (target), center (ingredient) - closest (target), center (ingredient) - kernel (target), center (ingredient) - separation (target) and center (ingredient) - shortest (target)) have a better discrimination performance with mean AUROC of 0.80 and mean AUPRC of 0.83, in contrast to that of the other models (**Figure 5**). Different from using the other distance metric at the ingredient level, the center-based models involve the identification of the central ingredients that have a minimal sum of shortest path lengths in the herb-ingredient network. The superior performance of the center-based distance models therefore suggests that the herb-pair interactions are mainly driven by few ingredients as determined as the center nodes. These topologically important ingredients may hold the key for understanding herb pair interactions.

**Figure 5:**
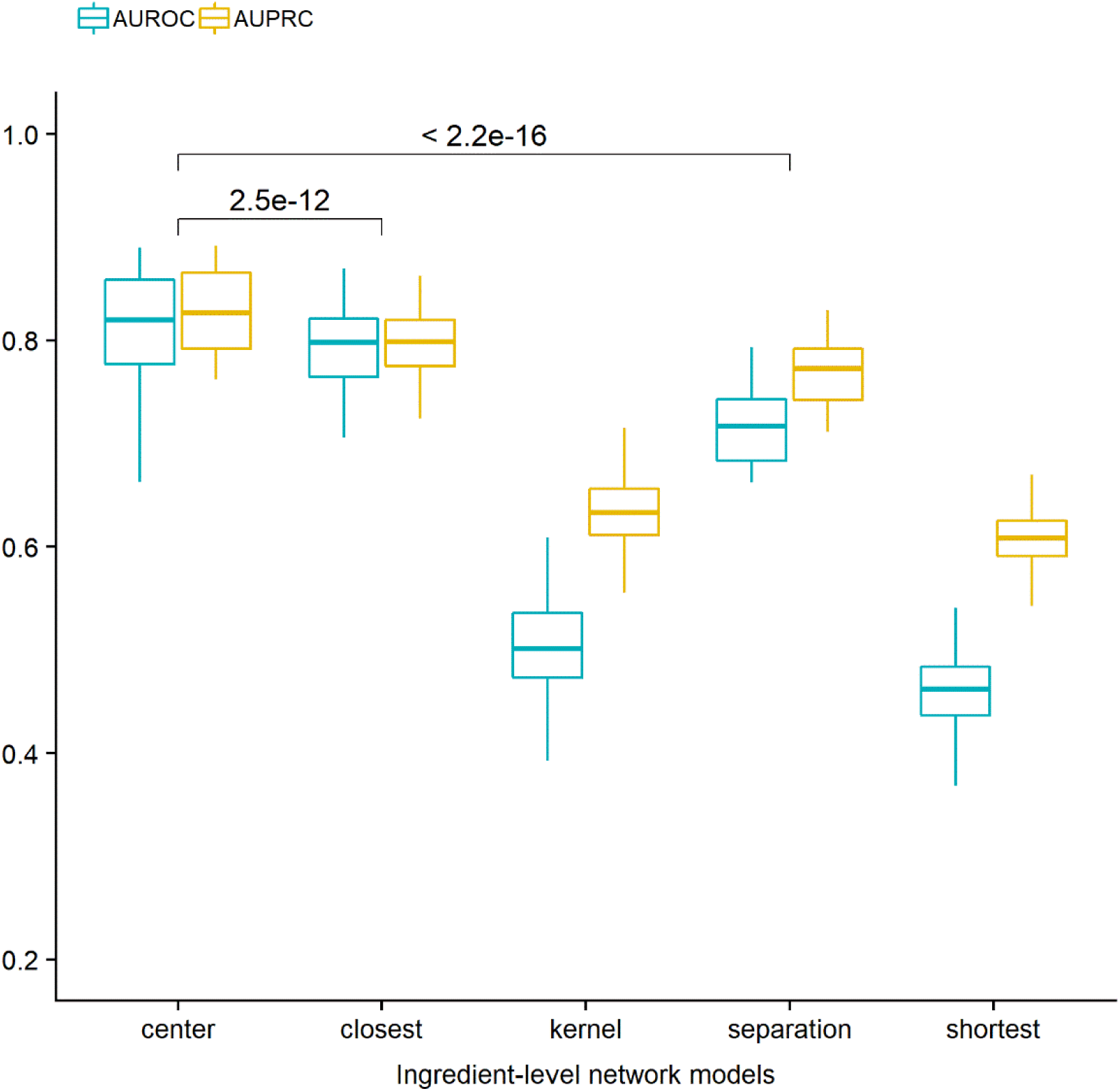
AUROC and AUPRC grouped by the distance models at the ingredient level. The statistical significance is determined by t-test.

To validate our hypothesis, we also collected 268 known herb pairs from the literature (**Supplementary Table 3**). We applied the 25 network models to evaluate how well these 268 known herb pairs can be separated from random pairs. In line with the previous results, we found that the distance between these known herb pairs is on average smaller than random pairs (**Supplementary Table 4**). The average AUROC and AUPRC across all the 25 models is 0.62 and 0.65, respectively. Furthermore, the center (ingredient) - shortest (target) model can achieve the top accuracy of AUROC 0.75 and AUPRC 0.73 (**Supplementary Table 4, Supplementary Figure 5**). Notably, the 268 known herb pairs were extracted from the literature that was independent from the datasets extracted from the TCMID. The overlap between these two datasets is minimal (n = 32), suggesting a general validity of using network models to predict the potential of herb pairs in TCM.

### 3.4 The combination mechanism of herb pair *Astragalus membranaceus* and *Glycyrrhiza uralensis*

We applied our network pharmacology modeling to the study of herb pair *Astragalus membranaceus* and *Glycyrrhiza uralensis*. The combination of *Astragalus membranaceus* and *Glycyrrhiza uralensis* has shown clinical efficacy to treat liver diseases by the inhibition of notch signaling pathways^65^. It was also reported that this herb pair is able to inhibit bile acid-stimulated inflammation in chronic cholestatic liver injury mice^56^ based on transcriptomics profiling^55^. However, their active ingredients and the mechanisms of action of remain poorly understood.

We retrieved 15 ingredients for *Astragalus membranaceus* and 27 ingredients for *Glycyrrhiza uralensis*, separately, for which three ingredients were common including *formononetin, clionasterol* and *clionasterol* (**Supplementary Table 5**). Based on the conclusion that center-based distance models tend to achieve better performance, we considered the distance for the herb pair as the distance of their center ingredients, which can be determined by five different models at the target level. We compared the herb distances with that of the top 200 herb pairs as well as the random herb pairs. We found that the herb pair distances are much smaller than that of the random herb pairs, suggesting a strong evidence for the close network proximity of the two herbs (**Table 2**).

**Table 2:**
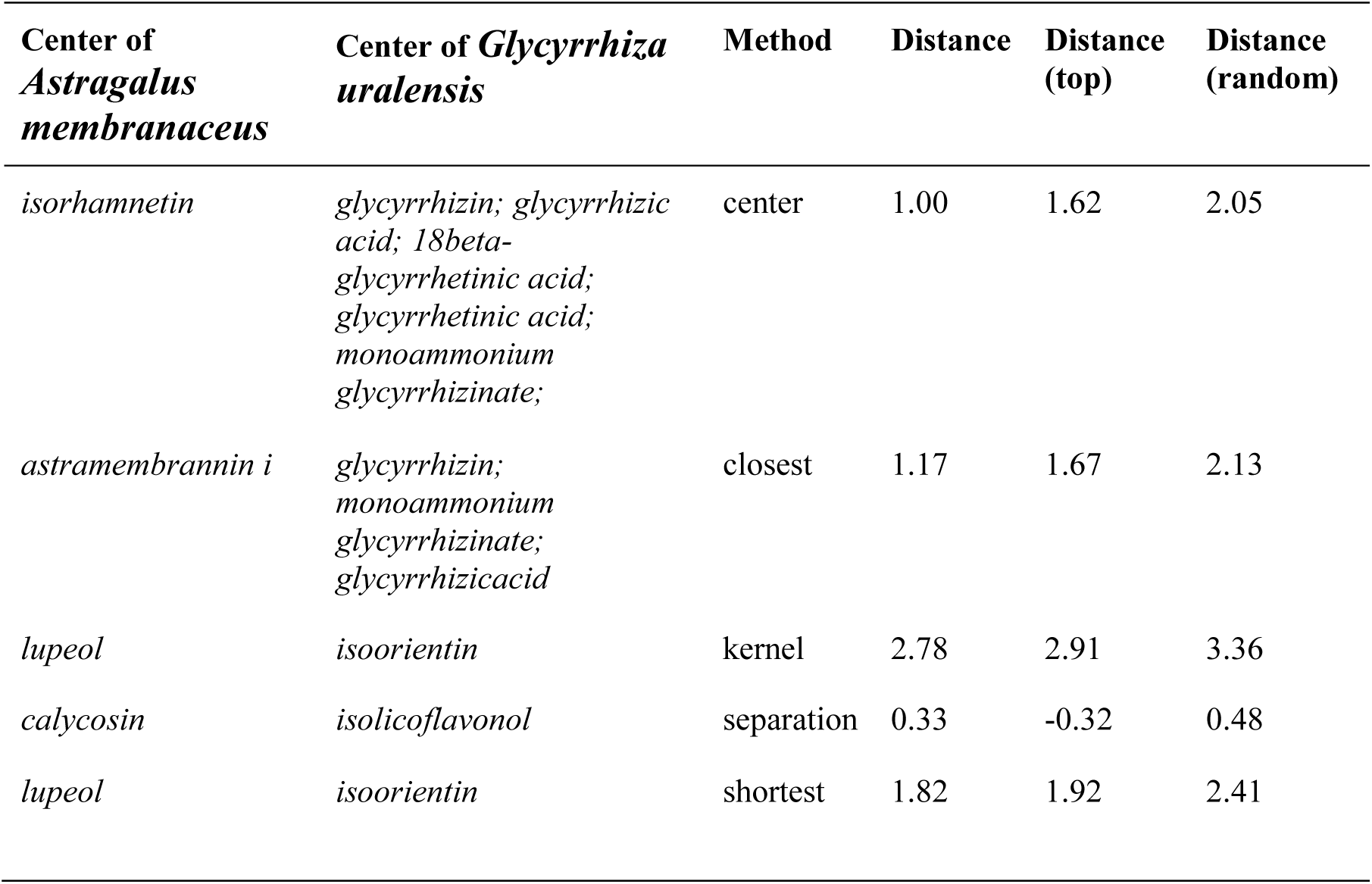
The center ingredients for *Astragalus membranaceus* and *Glycyrrhiza uralensis* determined by models with different distance methods at the target level while fixing the center distance method at the ingredient level.

| Center of *Astragalus membranaceus* | Center of *Glycyrrhiza uralensis* | Method | Distance | Distance (top) | Distance (random) |
| --- | --- | --- | --- | --- | --- |
| *isorhamnetin* | *glycyrrhizin*; *glycyrrhizic acid*; *18beta-glycyrrhetinic acid*; *glycyrrhetinic acid*; *monoammonium glycyrrhizinate*; | center | 1.00 | 1.62 | 2.05 |
| *astramembrannin i* | *glycyrrhizin*; *monoammonium glycyrrhizinate*; *glycyrrhizic acid* | closest | 1.17 | 1.67 | 2.13 |
| *lupeol* | *isoorientin* | kernel | 2.78 | 2.91 | 3.36 |
| *calycosin* | *isolicoflavonol* | separation | 0.33 | -0.32 | 0.48 |
| *lupeol* | *isoorientin* | shortest | 1.82 | 1.92 | 2.41 |

By applying the center (ingredient) - closest (target) model, we found that *astramembrannin i* and *glycyrrhizin* were identified as the center of *Astragalus membranaceus* and *Glycyrrhiza uralensis*, separately. It was shown that *glycyrrhizin* from *Glycyrrhiza uralensis* is effective on ferroptosis by inhibiting oxidative stress during acute liver failure^66^. Interestingly, it was reported that the synergistic anti-liver fibrosis actions by the *Astragalus membranaceus* and *Glycyrrhiza uralensis* can be attributed to the ingredient *astragalus saponins* from *Astragalus membranaceus* and ingredient of *glycyrrhizic acid* of *Glycyrrhiza uralensis* via TGF-β1/Smads signaling pathway modulation^67^, which is consistent with our analysis.

On the other hand, to apply the center (ingredient) - shortest (target) model, we first determined the shortest distance for each ingredient pair using the target interaction network, with which we can determine *lupeol* and *isoorientin* as the central ingredients of *Astragalus membranaceus* and *Glycyrrhiza uralensis*, separately. We found that the distance is 1.82, which is lower than the average (1.92) of the top herb pairs, and much lower than the average (2.41) of the random herb pairs. Interestingly, we found that the same center ingredients were also identified by the center (ingredient) - kernel (target) model. It was reported that *isoorientin* might protect alcohol induced hepatic fibrosis in rats by reducing the levels of inflammation-related pathways^68^. On the other hand, *lupeol* was known for protecting oxidative stress-induced cellular injury of mouse liver by downregulating anti-apoptotic Bcl-2 and upregulating pro-apoptotic Bax and Caspase 3^69^. To illustrate further the potential combinational effects of *lupeol* and *isoorientin*, we performed pathway analysis by the targets of these two ingredients (NFE2L2, AKT1 form *isoorientin* and CTNNB1, MITF, LSS, PTEN and TP53 form *lupeol*) (**Figure 6**). We found that these target genes are associated with pathways related to liver disease, especially the cholesterol biosynthesis pathway, the hepatocellular carcinoma pathway, the IL-5 signaling pathway as well as the ethanol metabolism resulting in production of ROS by the CYP2E1 pathway. Therefore, it is plausible that the anti-liver fibrosis effects of herb pair *Astragalus membranaceus* and *Glycyrrhiza uralensis* can be attributed to the combination of *lupeol* and *isoorientin*. Taken together, this case study exemplified the feasibility and rational of applying the network model to pinpoint potential ingredient interactions and their mechanisms of action.

**Figure 6:**
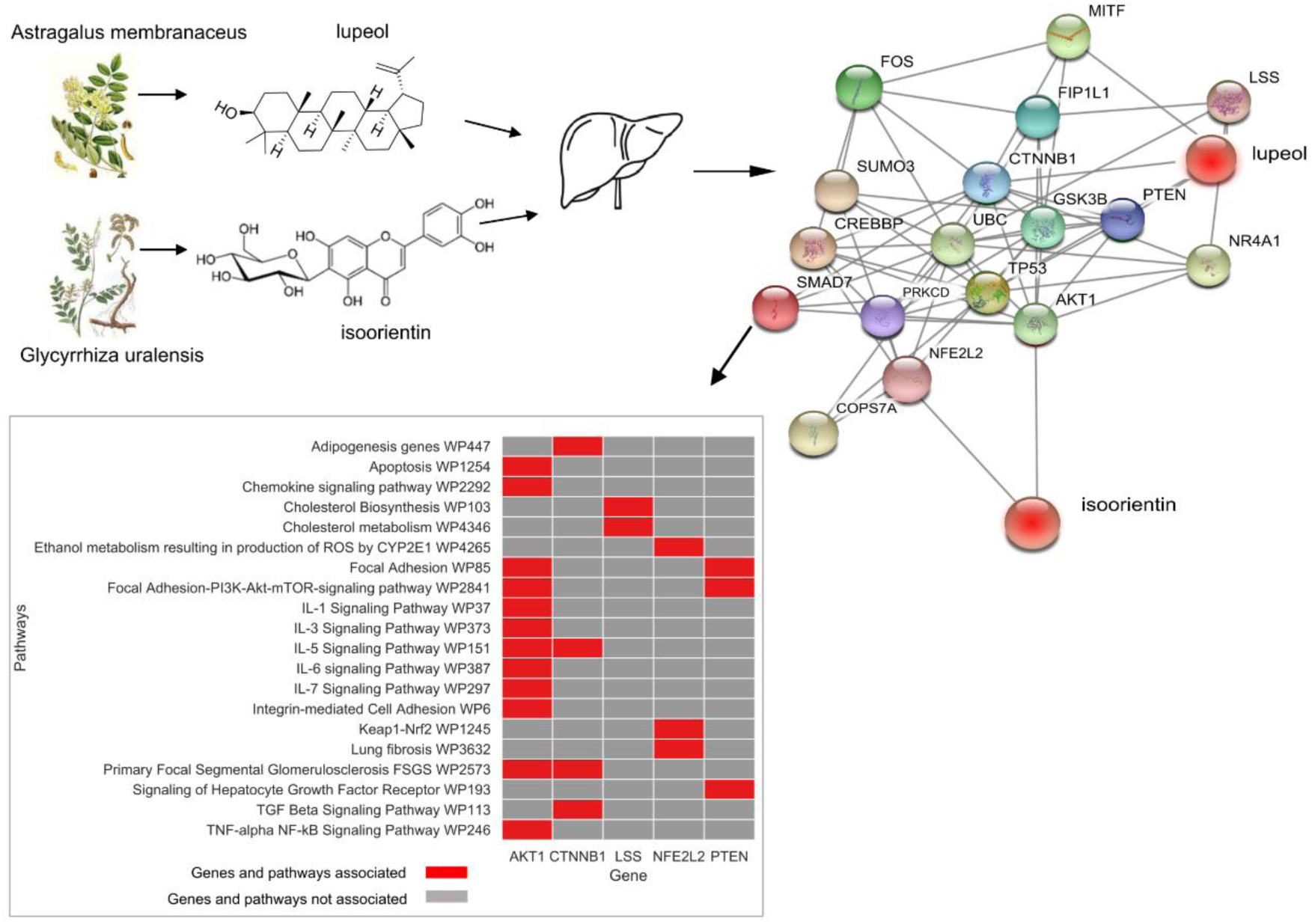
PPI network and pathway enrichment of the combination of *isoorientin* of *Glycyrrhiza uralensis* and *lupeol* of *Astragalus membranaceus*. The targets of the two center ingredients and their associated pathways are listed.

## 4 Discussion

Understanding the mechanisms of actions of TCM requires a more systematic investigation of the herb interactions. In this paper, we proposed a novel PPI-based network model to characterize the interaction of herb pairs. To illustrate the complex nature of TCM pharmacology, we developed network distance metrics by integrating the relationships between herb, ingredients and targets. We defined the herb-herb distance based on a multiple partite network which is commonly used for biological network modeling^35^. The components of such a multi-modal network include bipartite networks of herb-ingredient and ingredient-target interactions. We considered the network proximity distance at two levels, where the nodes of the networks can be either ingredients or targets. The two-level network modeling allows the characterization of herb-herb and ingredient-ingredient interactions with greater flexibility. In this study, we have provided a panel of 25 distance models, based on which we achieved a comprehensive evaluation of herb-herb interactions. Compared to the existing methods that are mainly focusing on single herbs, our network modeling can provide more insights on the mechanisms of action of TCM herb formulae, which by principle mainly involve multi-herb combinations.

We found that commonly used herb pairs tend to have smaller network proximity distance, suggesting stronger PPI interactions between them. Moreover, using the center distance at the ingredient level, the network model tends to achieve higher accuracy of discriminating the commonly used herb pairs from random herb pairs with the best AUROC of 0.87 and AUPRC of 0.87. In general, we found that the center distance at the ingredient level improved the prediction accuracy, suggesting that ingredients that are located in the center of the herb PPI network play important roles when combined with the other herbs. These center ingredients showed a minimal sum of shortest path lengths within the herb PPI network, and therefore are more likely to activate a cascade of multiple pathways. Prioritization of these center ingredients for further functional studies shall help us understand the synergistic effects of herb pairs. Using the herb pair *Astragalus membranaceus* and *Glycyrrhiza uralensis* as a case study, we confirmed that its network distance was shorter than that of random herb pairs. More interestingly, the potential synergistic effects of the center ingredient *lupeol* from *Astragalus membranaceus* and the center ingredient *isoorientin* from *Glycyrrhiza uralensis* were supported by the literature^67-69^, which warrants more experimental validation.

On the other hand, the stronger network proximity distance between the TCM herb pairs might be due to the overlapping ingredients. Indeed, we found that 86 out of 200 top common herb pairs shared at least one common ingredient. However, using the 114 herb pairs that do not share any common ingredients, we retained the same level of top prediction accuracy (AUROC 0.75 and AUPRC 0.73). Therefore, the strong PPI interactions were largely attributed by functionally related ingredients that may share common or similar targets. For example, the ingredient *nodakenin* from herb *Notopterygium incisum* and ingredient *limonene* from herb *Angelica pubescens f. biserrata* have five common targets, including NOS1, NOS2, NOS3, POR and MTRR. Targeting the same disease proteins with multiple ingredients is in fact an important strategy of TCM formula, as it may achieve the same level of efficacy while lowering the side effects that are caused by the high doses of single ingredient^15^.

Previously, Li et al. have proposed a Distance-Based-mutual-Information (DMIM) approach^14^ to determine an interaction score between herb pairs based on their frequencies. Compared to DMIM, our method is based on the information at deeper molecular levels such as herb-ingredient, ingredient-target and target-target relationships, which shall provide a more refined characterization of herb-interactions. However, there are limitations in our study that need to be improved in the future. For example, despite the knowledge of existing ingredients in an herb, their actual concentrations are largely unknown. Therefore, the current model treats each ingredient equally, which might lead to certain bias. Moreover, we empirically determined the common herb pairs by their frequencies of occurrences in TCM formulae, which might be suboptimal. On the other hand, we did not filter the ingredients by oral bioavailability (OB) and drug-likeness (DL) in our study of herb combinations, as it is known that ingredients in TCM with low OB or DL values may still play active roles due to their superior pharmacological properties^70^. Another limitation is the lack of target information for certain ingredients. In our model, we discarded the herbs and ingredients without any target information, as their biological roles remain unclear. In the future, computational methods, such as the similarity ensemble approach (SEA)^71^, and experimental methods such as thermal proteomics profiling (TPP)^72^ can help the TCM research in the aspect of targeted discovery of herb ingredients.

In conclusion, TCM formulae provide important resource of drug combinations in natural products. In this study, we proposed a network-based model to understand the rational of herb pairs in TCM. By qualifying the distances between herb pairs based on herb-ingredient-target interactions, the network model can identify the potential synergistic ingredients for which the mechanisms of action can be further explored. The modelling strategy itself not only helps us explore the space of herb combinations more effectively, but also can be used for prioritizing synergistic compound interactions that shall facilitate the drug discovery from TCM.

