## Supplementary for "Network-based modeling of herb combinations in Traditional Chinese Medicine"

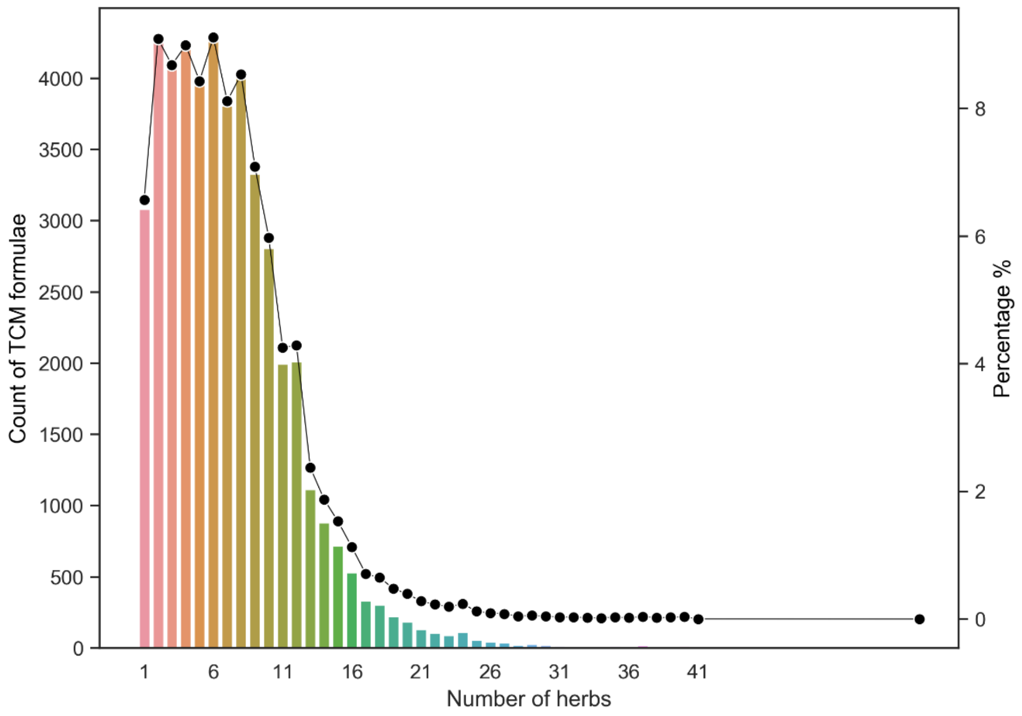


**Supplementary Figure 1. Distribution of the number of herbs in a TCM formula.** The left y-axis is the number of formulas with a specific number of herbs. The right y-axis is the percentage of the formulae with a specific number of herbs divided by the whole number of formulae. Most of the herb formulae (97.9 %) contain less than 20 herbs, with an average of 4.93.

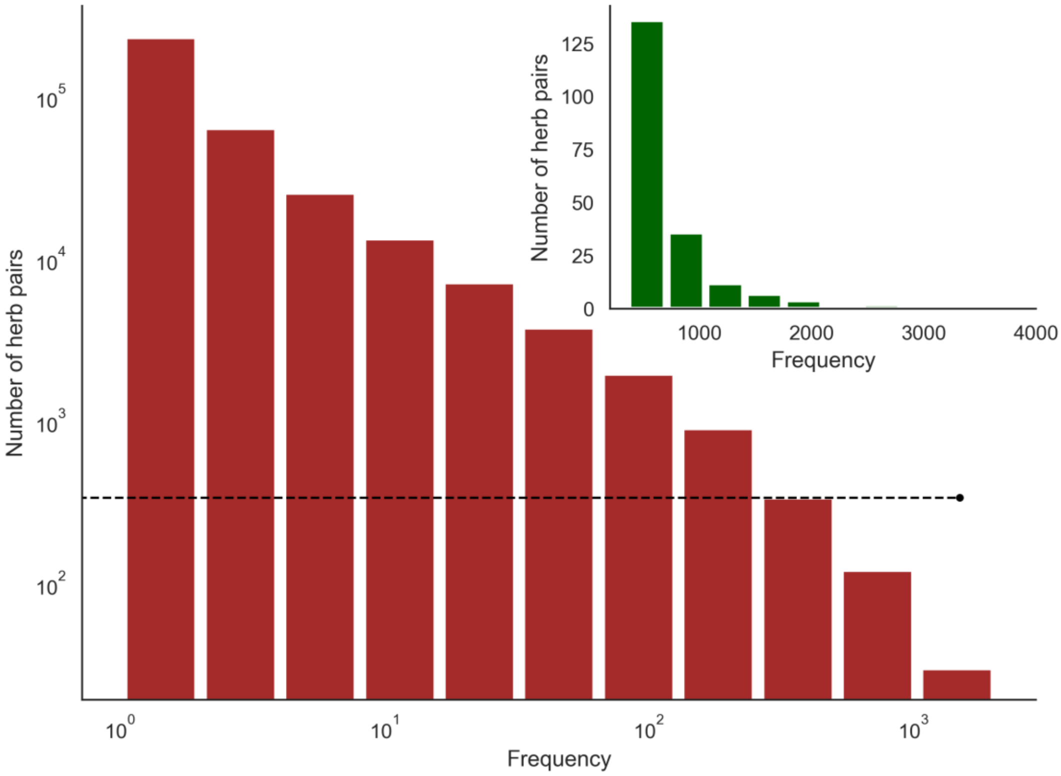


**Supplementary Figure 2. Distribution of herb pair frequency in the TCM formula.** The main figure shows the distribution of all herb pairs, while the top-right inset shows the distribution of herb pair frequency of top 200 herb pairs. It can be seen that the majority of the 349,197 herb pairs (99.4%) occurred in less than 100 herbal formulae. Only 163 herb pairs of the remaining 1950 (0.6%) herb pairs showed a frequency higher than 500. There is a sharp decrease of herb pair frequency after 200. Therefore, we considered those herb pairs with frequency larger than 200 to be the popular herb pairs.

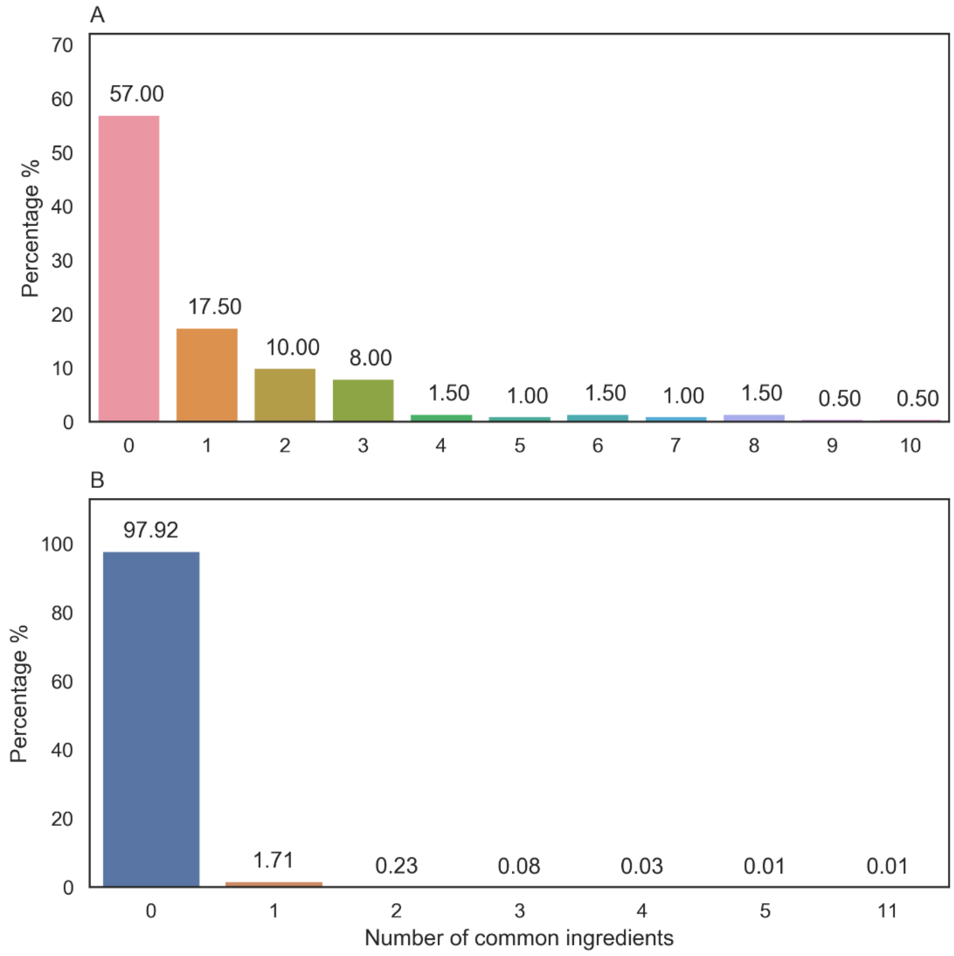


**Supplementary Figure 3. Distribution of the number of common ingredients for the top 200 herb pairs (top panel) as compared to randomly generated herb pairs (bottom panel).** These herb pairs involve 61 unique herbs, for which the average number of ingredients is 16.8. There is at least one common ingredient for 43% (86) of the top 200 herb pairs, while only 2.08% of randomly-generated herb pairs share at least one ingredient. There are 114 out of the 200 herb pairs which did not share any common ingredients, while a few herb pairs (n = 15) shared more than three ingredients.

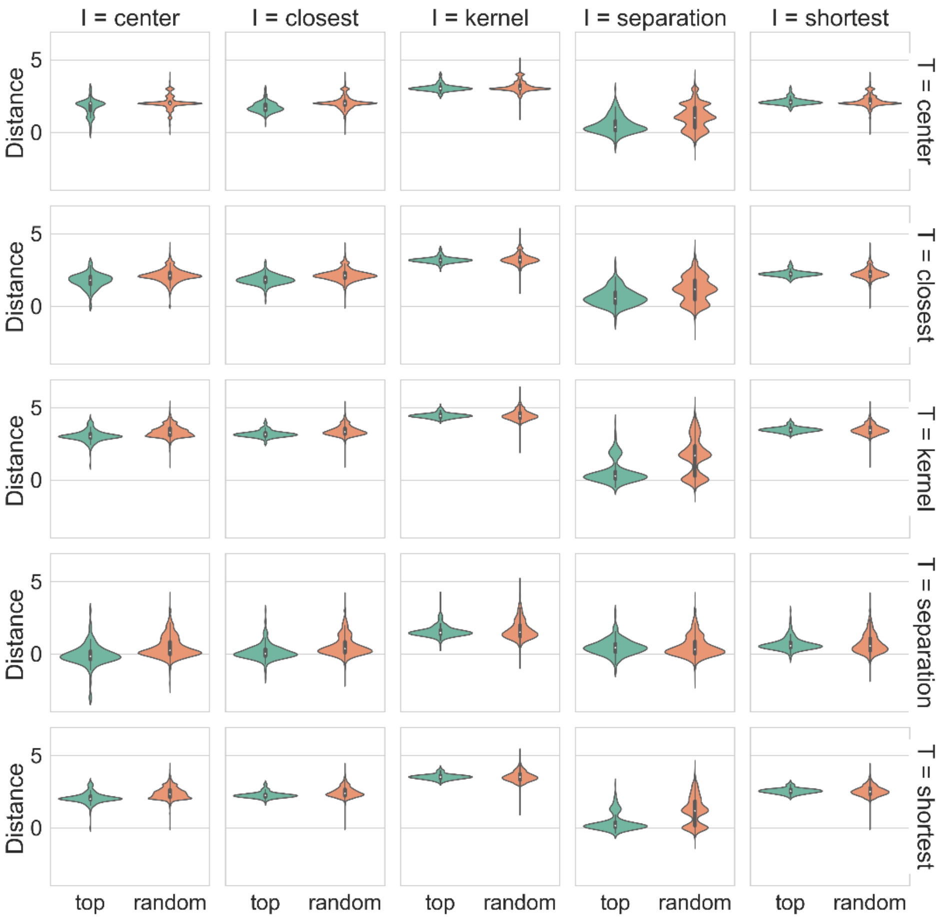


**Supplementary Figure 4. Network distances for top herb pairs and random herb pairs that share no common ingredients.** 'I' stands for the ingredient-level distance methods and 'T' stands for the target-level distance methods. Their distances are significantly lower than that for random herb pairs when we considered the 114 herb pairs that did not share any common ingredients, suggesting that target interactions from different ingredients remain a major strategy to affect functionally related pathways in a TCM herb pair.

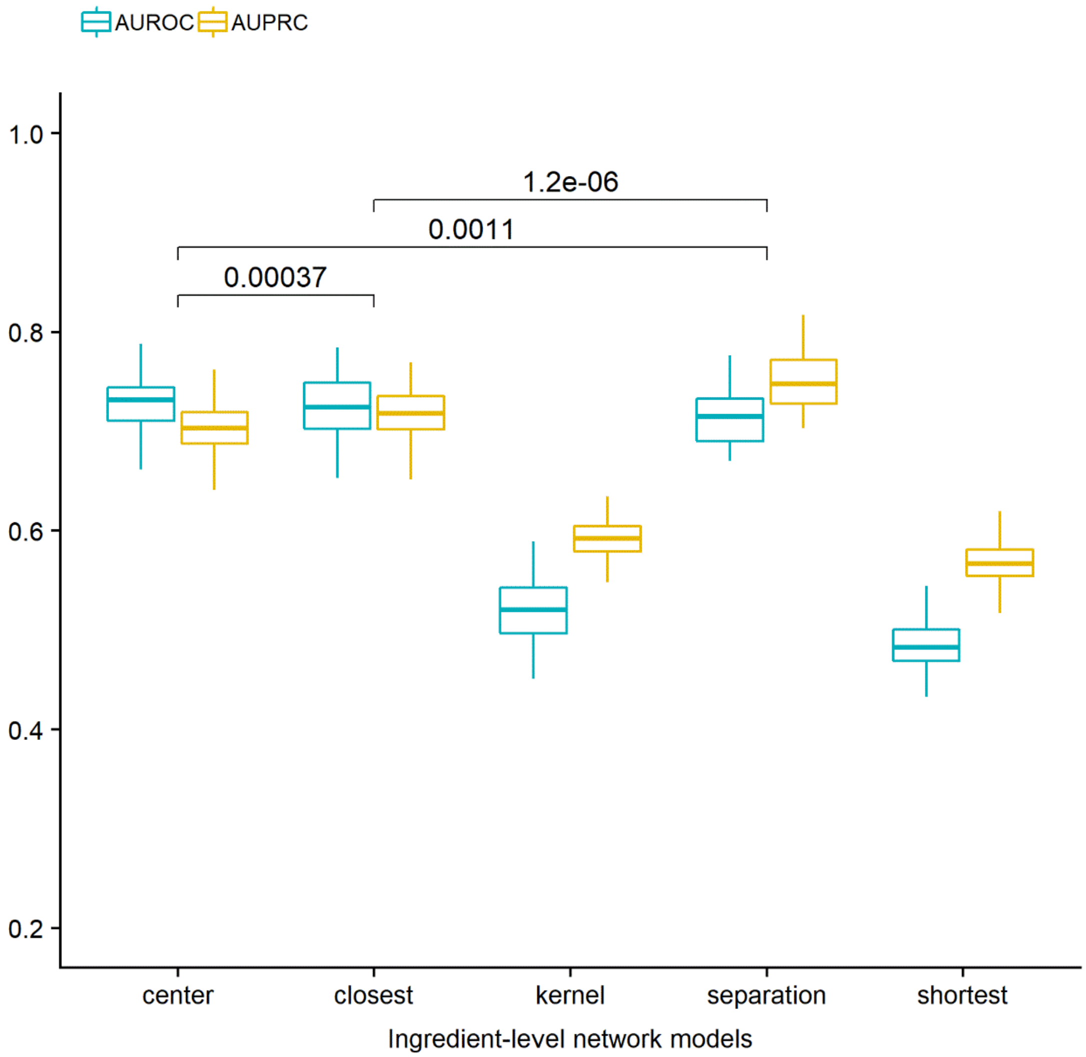


**Supplementary Figure 5. AUROC and AUPRC grouped by the distance models at the ingredient level for the literature-curated herb pairs.** The distance between these known herb pairs is on average smaller than random pairs, suggesting that the network models can separate these 268 known herb pairs from random pairs. The average AUROC and AUPRC is 0.62 and 0.65, respectively. Furthermore, the center (ingredient) - shortest (target) model can achieve the top accuracy of AUROC 0.75 and AUPRC 0.73.

### Supplementary Table 1. The most frequent herbs.

| Herb | Frequency | Chinese Name | English Name | Latin Name | Meridians | Pinyin Name | Properties | Used Part | TCMID herb id | TCMID herb website |
|---|---|---|---|---|---|---|---|---|---|---|
| GANCAO | 12518 | 甘草 | Ural Licorice Equivalent plant: Glycyrrhiza inflata, Glycyrrhiza glabra, Glycyrrhiza kansuensis, Glycyrrhiza aspera, Glycyrrhiza yunnanensis , Glycyrrhiza squamulosa | Glycyrrhiza uralensis | Lung,Spleen,Stomach,Heart | GANCAO | Mild,Sweet | root and rhizome | 6801 | http://119.3.41.228:8000/tcmid/herb/6801/ |
| DANGGUI | 7417 | 当归 | Chinese Angelica Equivalent plant: Phlojodicarpus sibiricus | Angelica sinensis | Spleen,Liver,Heart | DANGGUI | Warm,Pungent,Sweet | root | 2538 | http://119.3.41.228:8000/tcmid/herb/2538/ |
| RENSHEN | 7390 | 人参 | Ginseng | Panax ginseng [Syn. Panax schinseng] | Lung,Spleen,Heart | RENSHEN | Minor Warm,Sweet,Slightly Bitter | root | 3861 | http://119.3.41.228:8000/tcmid/herb/3861/ |

| | | | | | | | | | | |
|---|---|---|---|---|---|---|---|---|---|---|
| BAIZHU | 5259 | 白术 | Largehead Atractylodes | Atractylodes macrocephala [Syn. Atractylis macrocephala ] | Spleen,Stomach | BAIZHU | Warm,Sweet,Bitter | root | 7301 | http://119.3.41.228:8000/tcmid/herb/7301/ |
| HUANG QIN | 4163 | 黄芩 | Baikal Skullcap Equivalent plant: Scutellaria amoena , Scutellaria viscidula, Scutellaria likiangensis , Scutellaria rehderiana, Scutellaria hypericifolia | Scutellaria baicalensis | Lung,Large Intestine,Stomach,Small Intestine,Gallbladder | HUANG QIN | Cold,Bitter | root | 6700 | http://119.3.41.228:8000/tcmid/herb/6700/ |
| FANGFENG | 4074 | 防风 | Divaricate Saposhnikovia | Saposhnikovia divaricata [Syn. Ledebouriella seseloides] | Bladder,Spleen, Liver | FANGFENG | Minor Warm,Pungent,Sweet | root | 7847 | http://119.3.41.228:8000/tcmid/herb/7847/ |
| CHUAN XIONG | 4007 | 川芎 | Chuanxiong (Wallich Ligusticum) Equivalent plant: Cnidium officinale | Ligusticum chuanxiong [Syn. Ligusticum wallichii ] | Liver,Cardiovascular,Gallbladder | CHUAN XIONG | Warm,Pungent | rhizome | 5926 | http://119.3.41.228:8000/tcmid/herb/5926/ |

| | | | | | | | | | | |
|---|---|---|---|---|---|---|---|---|---|---|
| FU LING | 3666 | 茯苓 | Indian Bread | Poria cocos | Spleen,Heart,Kidney | FU LING | Mild,Sweet,Neutral | sclerotium | 7870 | http://119.3.41.228:8000/tcmid/herb/7870/ |
| CHEN PI | 3650 | 陈皮 | Dried Tangerine Peel | Pericarpium Citri Reticulatae | Lung,Spleen | CHEN PI | Warm,Pungent,Bitter | dried peel | 1849 | http://119.3.41.228:8000/tcmid/herb/1849/ |

### Supplementary Table 2. The most frequent herb pairs.

| TCMID Herb_1 id | Herb_1 Pinyin name | TCMID Herb_2 id | Herb_2 Pinyin name | Frequency | Herb_1 Latin name | Herb_2 Latin name |
|---|---|---|---|---|---|---|
| 3861 | REN SHEN | 6801 | GAN CAO | 3846 | Panax ginseng | Glycyrrhiza uralensis |
| 2538 | DANG GUI | 6801 | GAN CAO | 2907 | Angelica sinensis | Glycyrrhiza uralensis |
| 3861 | REN SHEN | 7301 | BAI ZHU | 2599 | Panax ginseng | Atractylodes macrocephala |
| 6801 | GAN CAO | 7301 | BAI ZHU | 2578 | Glycyrrhiza uralensis | Atractylodes macrocephala |
| 6700 | HUANG QIN | 6801 | GAN CAO | 2265 | Scutellaria baicalensis | Glycyrrhiza uralensis |
| 2538 | DANG GUI | 3861 | REN SHEN | 2097 | Angelica sinensis | Panax ginseng |
| 1849 | CHEN PI | 6801 | GAN CAO | 2002 | Pericarpium Citri Reticulatae | Glycyrrhiza uralensis |
| 6801 | GAN CAO | 7847 | FANG FENG | 1960 | Glycyrrhiza uralensis | Saposhnikovia divaricata |
| 6801 | GAN CAO | 7870 | FU LING | 1817 | Glycyrrhiza uralensis | Poria cocos |
| 2538 | DANG GUI | 7301 | BAI ZHU | 1716 | Angelica sinensis | Atractylodes macrocephala |
| 5800 | BAN XIA | 6801 | GAN CAO | 1689 | Pinellia ternata | Glycyrrhiza uralensis |
| 3507 | JIE GENG | 6801 | GAN CAO | 1665 | Platycodon grandiflorum | Glycyrrhiza uralensis |
| 3861 | REN SHEN | 7870 | FU LING | 1508 | Panax ginseng | Poria cocos |
| 3861 | REN SHEN | 6919 | HUANG QI | 1506 | Panax ginseng | Astragalus membranaceus |
| 7301 | BAI ZHU | 7870 | FU LING | 1480 | Atractylodes macrocephala | Poria cocos |

| | | | | | | |
|---|---|---|---|---|---|---|
| 6801 | GAN CAO | 6919 | HUANG QI | 1411 | Glycyrrhiza uralensis | Astragalus membranaceus |
| 2538 | DANG GUI | 3997 | BAI SHAO | 1328 | Angelica sinensis | Paeonia albiflora |
| 1849 | CHEN PI | 7301 | BAI ZHU | 1305 | Pericarpium Citri Reticulatae | Atractylodes macrocephala |
| 5777 | ZHI KE | 6801 | GAN CAO | 1299 | Citrus aurantium | Glycyrrhiza uralensis |
| 4776 | QIANG HUO | 7847 | FANG FENG | 1292 | Notopterygium incisum | Saposhnikovia divaricata |
| 2538 | DANG GUI | 6919 | HUANG QI | 1258 | Angelica sinensis | Astragalus membranaceus |
| 6801 | GAN CAO | 7648 | HUANG LIAN | 1217 | Glycyrrhiza uralensis | Coptis chinensis |
| 2538 | DANG GUI | 7847 | FANG FENG | 1169 | Angelica sinensis | Saposhnikovia divaricata |
| 3867 | HOU PO | 6801 | GAN CAO | 1155 | Magnolia officinalis | Glycyrrhiza uralensis |
| 1656 | GAN JIANG | 6801 | GAN CAO | 1126 | Zingiber officinale | Glycyrrhiza uralensis |
| 3861 | REN SHEN | 5800 | BAN XIA | 1113 | Panax ginseng | Pinellia ternata |
| 5151 | MU XIANG | 6801 | GAN CAO | 1079 | Saussurea lappa | Glycyrrhiza uralensis |
| 6700 | HUANG QIN | 7648 | HUANG LIAN | 1075 | Scutellaria baicalensis | Coptis chinensis |
| 1849 | CHEN PI | 3861 | REN SHEN | 1048 | Pericarpium Citri Reticulatae | Panax ginseng |
| 2538 | DANG GUI | 7870 | FU LING | 1006 | Angelica sinensis | Poria cocos |
| 3997 | BAI SHAO | 6801 | GAN CAO | 1004 | Paeonia albiflora | Glycyrrhiza uralensis |
| 1849 | CHEN PI | 2538 | DANG GUI | 1003 | Pericarpium Citri Reticulatae | Angelica sinensis |
| 1849 | CHEN PI | 7870 | FU LING | 976 | Pericarpium Citri Reticulatae | Poria cocos |
| 4776 | QIANG HUO | 6801 | GAN CAO | 973 | Notopterygium incisum | Glycyrrhiza uralensis |
| 2538 | DANG GUI | 6700 | HUANG QIN | 959 | Angelica sinensis | Scutellaria baicalensis |
| 1849 | CHEN PI | 5800 | BAN XIA | 944 | Pericarpium Citri Reticulatae | Pinellia ternata |
| 3861 | REN SHEN | 7847 | FANG FENG | 941 | Panax ginseng | Saposhnikovia divaricata |

| | | | | | | |
|---|---|---|---|---|---|---|
| 6919 | HUANG QI | 7301 | BAI ZHU | 926 | Astragalus membranaceus | Atractylodes macrocephala |
| 1660 | MA HUANG | 6801 | GAN CAO | 925 | Ephedra sinica | Glycyrrhiza uralensis |
| 5800 | BAN XIA | 7301 | BAI ZHU | 880 | Pinellia ternata | Atractylodes macrocephala |
| 3861 | REN SHEN | 5151 | MU XIANG | 871 | Panax ginseng | Saussurea lappa |
| 6801 | GAN CAO | 7047 | DA HUANG | 869 | Glycyrrhiza uralensis | Rheum officinale |
| 6801 | GAN CAO | 7419 | XING REN | 864 | Glycyrrhiza uralensis | Prunus armeniaca |
| 1656 | GAN JIANG | 3861 | REN SHEN | 856 | Zingiber officinale | Panax ginseng |
| 2538 | DANG GUI | 5151 | MU XIANG | 844 | Angelica sinensis | Saussurea lappa |
| 6700 | HUANG QIN | 7847 | FANG FENG | 825 | Scutellaria baicalensis | Saposhnikovia divaricata |
| 5151 | MU XIANG | 7301 | BAI ZHU | 818 | Saussurea lappa | Atractylodes macrocephala |
| 6700 | HUANG QIN | 7047 | DA HUANG | 810 | Scutellaria baicalensis | Rheum officinale |
| 2538 | DANG GUI | 7648 | HUANG LIAN | 808 | Angelica sinensis | Coptis chinensis |
| 5800 | BAN XIA | 7870 | FU LING | 807 | Pinellia ternata | Poria cocos |
| 5107 | SHI GAO | 6801 | GAN CAO | 786 | Gypsum Fibrosum | Glycyrrhiza uralensis |
| 1656 | GAN JIANG | 7301 | BAI ZHU | 784 | Zingiber officinale | Atractylodes macrocephala |
| 3006 | SHENG MA | 6801 | GAN CAO | 778 | Cimicifuga foetida | Glycyrrhiza uralensis |
| 4526 | JING JIE | 7847 | FANG FENG | 775 | Schizonepeta tenuifolia | Saposhnikovia divaricata |
| 3867 | HOU PO | 7301 | BAI ZHU | 752 | Magnolia officinalis | Atractylodes macrocephala |
| 3997 | BAI SHAO | 7301 | BAI ZHU | 752 | Paeonia albiflora | Atractylodes macrocephala |
| 4518 | LIAN QIAO | 6801 | GAN CAO | 748 | Forsythia suspensa | Glycyrrhiza uralensis |
| 3861 | REN SHEN | 6700 | HUANG QIN | 742 | Panax ginseng | Scutellaria baicalensis |
| 5258 | CANG ZHU | 6801 | GAN CAO | 742 | Atractylodes lancea | Glycyrrhiza uralensis |
| 1173 | FU ZI | 1656 | GAN JIANG | 736 | Aconitum carmichaeli | Zingiber officinale |

| | | | | | | |
|---|---|---|---|---|---|---|
| 1849 | CHEN PI | 3867 | HOU PO | 722 | Pericarpium Citri Reticulatae | Magnolia officinalis |
| 3507 | JIE GENG | 3861 | REN SHEN | 717 | Platycodon grandiflorum | Panax ginseng |
| 3861 | REN SHEN | 3867 | HOU PO | 714 | Panax ginseng | Magnolia officinalis |
| 1304 | ZHI MU | 6801 | GAN CAO | 713 | Anemarrhena asphodeloides | Glycyrrhiza uralensis |
| 3507 | JIE GENG | 7847 | FANG FENG | 702 | Platycodon grandiflorum | Saposhnikovia divaricata |
| 1849 | CHEN PI | 5151 | MU XIANG | 701 | Pericarpium Citri Reticulatae | Saussurea lappa |
| 1173 | FU ZI | 3861 | REN SHEN | 687 | Aconitum carmichaeli | Panax ginseng |
| 2538 | DANG GUI | 4776 | QIANG HUO | 681 | Angelica sinensis | Notopterygium incisum |
| 1656 | GAN JIANG | 2538 | DANG GUI | 674 | Zingiber officinale | Angelica sinensis |
| 1518 | WU WEI ZI | 3861 | REN SHEN | 668 | Schisandra chinensis | Panax ginseng |
| 3656 | CHI FU LING | 6801 | GAN CAO | 661 | | Glycyrrhiza uralensis |
| 1173 | FU ZI | 6801 | GAN CAO | 655 | Aconitum carmichaeli | Glycyrrhiza uralensis |
| 3507 | JIE GENG | 6700 | HUANG QIN | 653 | Platycodon grandiflorum | Scutellaria baicalensis |
| 2538 | DANG GUI | 5777 | ZHI KE | 649 | Angelica sinensis | Citrus aurantium |
| 6338 | QIAN HU | 6801 | GAN CAO | 649 | Angelica decursiva | Glycyrrhiza uralensis |
| 4526 | JING JIE | 6801 | GAN CAO | 648 | Schizonepeta tenuifolia | Glycyrrhiza uralensis |
| 1849 | CHEN PI | 6196 | QING PI | 643 | Pericarpium Citri Reticulatae | Pericarpium Citri Reticulatae Viride |
| 2276 | MU TONG | 6801 | GAN CAO | 642 | Akebia quinata | Glycyrrhiza uralensis |
| 3861 | REN SHEN | 3997 | BAI SHAO | 627 | Panax ginseng | Paeonia albiflora |
| 1010 | YUAN ZHI | 3861 | REN SHEN | 621 | Polygala tenuifolia | Panax ginseng |
| 2538 | DANG GUI | 7047 | DA HUANG | 621 | Angelica sinensis | Rheum officinale |

| | | | | | | |
|---|---|---|---|---|---|---|
| 7648 | HUANG LIAN | 8114 | HUANG BAI | 617 | Coptis chinensis | Phellodendron amurense |
| 2538 | DANG GUI | 3507 | JIE GENG | 604 | Angelica sinensis | Platycodon grandiflorum |
| 1849 | CHEN PI | 5258 | CANG ZHU | 604 | Pericarpium Citri Reticulatae | Atractylodes lancea |
| 4449 | DU HUO | 7847 | FANG FENG | 602 | Angelica pubescens f. biserrata | Saposhnikovia divaricata |
| 1173 | FU ZI | 2538 | DANG GUI | 590 | Aconitum carmichaeli | Angelica sinensis |
| 1173 | FU ZI | 7301 | BAI ZHU | 589 | Aconitum carmichaeli | Atractylodes macrocephala |
| 3861 | REN SHEN | 5421 | FU SHEN | 587 | Panax ginseng | Poria |
| 4428 | SHE XIANG | 5260 | ZHU SHA | 579 | Moschus moschiferus, Moschus berezovskii, Moschus sifanicus | Cinnabaris |
| 3507 | JIE GENG | 5777 | ZHI KE | 570 | Platycodon grandiflorum | Citrus aurantium |
| 1849 | CHEN PI | 5777 | ZHI KE | 570 | Pericarpium Citri Reticulatae | Citrus aurantium |
| 2538 | DANG GUI | 5736 | NIU XI | 564 | Angelica sinensis | Achyranthes bidentata |
| 4518 | LIAN QIAO | 6700 | HUANG QIN | 564 | Forsythia suspensa | Scutellaria baicalensis |
| 7301 | BAI ZHU | 7847 | FANG FENG | 559 | Atractylodes macrocephala | Saposhnikovia divaricata |
| 6801 | GAN CAO | 7484 | SHENG JIANG | 557 | Glycyrrhiza uralensis | Zingiber officinale |
| 5151 | MU XIANG | 5374 | DING XIANG | 552 | Saussurea lappa | Syzygium aromaticum |
| 3867 | HOU PO | 5151 | MU XIANG | 548 | Magnolia officinalis | Saussurea lappa |
| 2538 | DANG GUI | 5800 | BAN XIA | 547 | Angelica sinensis | Pinellia ternata |
| 1849 | CHEN PI | 3997 | BAI SHAO | 546 | Pericarpium Citri Reticulatae | Paeonia albiflora |
| 3997 | BAI SHAO | 7870 | FU LING | 544 | Paeonia albiflora | Poria cocos |
| 3861 | REN SHEN | 5777 | ZHI KE | 539 | Panax ginseng | Citrus aurantium |

| | | | | | | |
|---|---|---|---|---|---|---|
| 2538 | DANG GUI | 3867 | HOU PO | 535 | Angelica sinensis | Magnolia officinalis |
| 1518 | WU WEI ZI | 6801 | GAN CAO | 535 | Schisandra chinensis | Glycyrrhiza uralensis |
| 6801 | GAN CAO | 7704 | CHI SHAO YAO | 532 | Glycyrrhiza uralensis | |
| 6700 | HUANG QIN | 7301 | BAI ZHU | 530 | Scutellaria baicalensis | Atractylodes macrocephala |
| 6919 | HUANG QI | 7847 | FANG FENG | 528 | Astragalus membranaceus | Saposhnikovia divaricata |
| 6196 | QING PI | 6801 | GAN CAO | 528 | Pericarpium Citri Reticulatae Viride | Glycyrrhiza uralensis |
| 4518 | LIAN QIAO | 7847 | FANG FENG | 528 | Forsythia suspensa | Saposhnikovia divaricata |
| 3867 | HOU PO | 5800 | BAN XIA | 527 | Magnolia officinalis | Pinellia ternata |
| 5777 | ZHI KE | 6700 | HUANG QIN | 523 | Citrus aurantium | Scutellaria baicalensis |
| 4449 | DU HUO | 4776 | QIANG HUO | 522 | Angelica pubescens f. biserrata | Notopterygium incisum |
| 1849 | CHEN PI | 7520 | XIANG FU | 522 | Pericarpium Citri Reticulatae | Cyperus rotundus |
| 5034 | XUAN SHEN | 6801 | GAN CAO | 521 | Scrophularia ningpoensis | Glycyrrhiza uralensis |
| 1849 | CHEN PI | 3507 | JIE GENG | 520 | Pericarpium Citri Reticulatae | Platycodon grandiflorum |
| 5800 | BAN XIA | 6700 | HUANG QIN | 517 | Pinellia ternata | Scutellaria baicalensis |
| 2538 | DANG GUI | 7520 | XIANG FU | 507 | Angelica sinensis | Cyperus rotundus |
| 6801 | GAN CAO | 7520 | XIANG FU | 506 | Glycyrrhiza uralensis | Cyperus rotundus |
| 5151 | MU XIANG | 5777 | ZHI KE | 505 | Saussurea lappa | Citrus aurantium |
| 5777 | ZHI KE | 7847 | FANG FENG | 503 | Citrus aurantium | Saposhnikovia divaricata |
| 3861 | REN SHEN | 7648 | HUANG LIAN | 495 | Panax ginseng | Coptis chinensis |
| 3379 | ZHI SHI | 6801 | GAN CAO | 487 | Citrus aurantium | Glycyrrhiza uralensis |
| 7047 | DA HUANG | 7648 | HUANG LIAN | 486 | Rheum officinale | Coptis chinensis |

| | | | | | | |
|---|---|---|---|---|---|---|
| 2538 | DANG GUI | 3884 | BAI SHAO YAO | 486 | Angelica sinensis | |
| 5151 | MU XIANG | 6726 | CHEN XIANG | 485 | Saussurea lappa | Aquilaria agallocha |
| 6205 | GE GEN | 6801 | GAN CAO | 484 | Pueraria lobata | Glycyrrhiza uralensis |
| 2538 | DANG GUI | 7704 | CHI SHAO YAO | 484 | Angelica sinensis | |
| 1849 | CHEN PI | 6700 | HUANG QIN | 483 | Pericarpium Citri Reticulatae | Scutellaria baicalensis |
| 5777 | ZHI KE | 5800 | BAN XIA | 477 | Citrus aurantium | Pinellia ternata |
| 4428 | SHE XIANG | 6238 | XIONG HUANG | 474 | Moschus moschiferus, Moschus berezovskii, Moschus sifanicus | Realgar |
| 1304 | ZHI MU | 6700 | HUANG QIN | 472 | Anemarrhena asphodeloides | Scutellaria baicalensis |
| 2522 | MAI DONG | 6801 | GAN CAO | 469 | Ophiopogon japonicus | Glycyrrhiza uralensis |
| 1656 | GAN JIANG | 3867 | HOU PO | 464 | Zingiber officinale | Magnolia officinalis |
| 2538 | DANG GUI | 7524 | ROU GUI | 463 | Angelica sinensis | Cinnamomum cassia |
| 5151 | MU XIANG | 6196 | QING PI | 462 | Saussurea lappa | Pericarpium Citri Reticulatae Viride |
| 6801 | GAN CAO | 8114 | HUANG BAI | 460 | Glycyrrhiza uralensis | Phellodendron amurense |
| 5421 | FU SHEN | 6801 | GAN CAO | 459 | Poria | Glycyrrhiza uralensis |
| 4449 | DU HUO | 6801 | GAN CAO | 459 | Angelica pubescens f. biserrata | Glycyrrhiza uralensis |
| 6700 | HUANG QIN | 8114 | HUANG BAI | 459 | Scutellaria baicalensis | Phellodendron amurense |
| 6700 | HUANG QIN | 7870 | FU LING | 450 | Scutellaria baicalensis | Poria cocos |
| 2538 | DANG GUI | 8114 | HUANG BAI | 449 | Angelica sinensis | Phellodendron amurense |
| 1660 | MA HUANG | 7419 | XING REN | 446 | Ephedra sinica | Prunus armeniaca |

| | | | | | | |
|---|---|---|---|---|---|---|
| 7648 | HUANG LIAN | 7847 | FANG FENG | 443 | Coptis chinensis | Saposhnikovia divaricata |
| 3884 | BAI SHAO YAO | 6801 | GAN CAO | 443 | | Glycyrrhiza uralensis |
| 3867 | HOU PO | 7870 | FU LING | 441 | Magnolia officinalis | Poria cocos |
| 6919 | HUANG QI | 7870 | FU LING | 438 | Astragalus membranaceus | Poria cocos |
| 3656 | CHI FU LING | 3861 | REN SHEN | 434 | | Panax ginseng |
| 3006 | SHENG MA | 6700 | HUANG QIN | 434 | Cimicifuga foetida | Scutellaria baicalensis |
| 3507 | JIE GENG | 7870 | FU LING | 434 | Platycodon grandiflorum | Poria cocos |
| 3507 | JIE GENG | 5800 | BAN XIA | 433 | Platycodon grandiflorum | Pinellia ternata |
| 3167 | BEI MU | 6801 | GAN CAO | 431 | Bulbus Fritillariae | Glycyrrhiza uralensis |
| 3507 | JIE GENG | 4518 | LIAN QIAO | 431 | Platycodon grandiflorum | Forsythia suspensa |
| 1304 | ZHI MU | 3861 | REN SHEN | 430 | Anemarrhena asphodeloides | Panax ginseng |
| 4776 | QIANG HUO | 6700 | HUANG QIN | 428 | Notopterygium incisum | Scutellaria baicalensis |
| 5106 | DI GU PI | 6801 | GAN CAO | 426 | Cortex Lycii Radicis | Glycyrrhiza uralensis |
| 5260 | ZHU SHA | 6238 | XIONG HUANG | 426 | Cinnabaris | Realgar |
| 3507 | JIE GENG | 7301 | BAI ZHU | 422 | Platycodon grandiflorum | Atractylodes macrocephala |
| 1660 | MA HUANG | 7847 | FANG FENG | 422 | Ephedra sinica | Saposhnikovia divaricata |
| 1656 | GAN JIANG | 5151 | MU XIANG | 417 | Zingiber officinale | Saussurea lappa |
| 1957 | CHI SHAO | 2538 | DANG GUI | 415 | Radix Paeoniae Rubra | Angelica sinensis |
| 2522 | MAI DONG | 3861 | REN SHEN | 414 | Ophiopogon japonicus | Panax ginseng |
| 3867 | HOU PO | 5258 | CANG ZHU | 412 | Magnolia officinalis | Atractylodes lancea |
| 2474 | HONG HUA | 2538 | DANG GUI | 411 | Carthamus tinctorius | Angelica sinensis |
| 1010 | YUAN ZHI | 6801 | GAN CAO | 410 | Polygala tenuifolia | Glycyrrhiza uralensis |

| | | | | | | |
|---|---|---|---|---|---|---|
| 5107 | SHI GAO | 6700 | HUANG QIN | 407 | Gypsum Fibrosum | Scutellaria baicalensis |
| 3861 | REN SHEN | 7524 | ROU GUI | 406 | Panax ginseng | Cinnamomum cassia |
| 1957 | CHI SHAO | 6801 | GAN CAO | 406 | Radix Paeoniae Rubra | Glycyrrhiza uralensis |
| 2538 | DANG GUI | 2834 | RU XIANG | 405 | Angelica sinensis | Boswellia carterii |
| 7301 | BAI ZHU | 7524 | ROU GUI | 404 | Atractylodes macrocephala | Cinnamomum cassia |
| 5258 | CANG ZHU | 7301 | BAI ZHU | 403 | Atractylodes lancea | Atractylodes macrocephala |
| 4428 | SHE XIANG | 4459 | NIU HUANG | 400 | Moschus moschiferus, Moschus berezovskii, Moschus sifanicus | Bos taurus domesticus, Bubalus bubalis |
| 5151 | MU XIANG | 5800 | BAN XIA | 400 | Saussurea lappa | Pinellia ternata |
| 2834 | RU XIANG | 4428 | SHE XIANG | 400 | Boswellia carterii | Moschus moschiferus, Moschus berezovskii, Moschus sifanicus |
| 1849 | CHEN PI | 7648 | HUANG LIAN | 398 | Pericarpium Citri Reticulatae | Coptis chinensis |
| 5777 | ZHI KE | 7301 | BAI ZHU | 397 | Citrus aurantium | Atractylodes macrocephala |
| 2538 | DANG GUI | 4518 | LIAN QIAO | 396 | Angelica sinensis | Forsythia suspensa |
| 5151 | MU XIANG | 7870 | FU LING | 396 | Saussurea lappa | Poria cocos |
| 2538 | DANG GUI | 4526 | JING JIE | 391 | Angelica sinensis | Schizonepeta tenuifolia |
| 6741 | TIAN MA | 7847 | FANG FENG | 390 | Gastrodia elata | Saposhnikovia divaricata |
| 7301 | BAI ZHU | 7648 | HUANG LIAN | 389 | Atractylodes macrocephala | Coptis chinensis |
| 3867 | HOU PO | 5777 | ZHI KE | 389 | Magnolia officinalis | Citrus aurantium |
| 7847 | FANG FENG | 7870 | FU LING | 388 | Saposhnikovia divaricata | Poria cocos |

| | | | | | | |
|---|---|---|---|---|---|---|
| 2538 | DANG GUI | 4449 | DU HUO | 386 | Angelica sinensis | Angelica pubescens f. biserrata |
| 5258 | CANG ZHU | 7847 | FANG FENG | 384 | Atractylodes lancea | Saposhnikovia divaricata |
| 3006 | SHENG MA | 7847 | FANG FENG | 383 | Cimicifuga foetida | Saposhnikovia divaricata |
| 6801 | GAN CAO | 7524 | ROU GUI | 382 | Glycyrrhiza uralensis | Cinnamomum cassia |
| 3997 | BAI SHAO | 6700 | HUANG QIN | 379 | Paeonia albiflora | Scutellaria baicalensis |
| 2538 | DANG GUI | 4256 | SHENG DI HUANG | 379 | Angelica sinensis | Radix Rehmanniae |
| 3861 | REN SHEN | 4776 | QIANG HUO | 373 | Panax ginseng | Notopterygium incisum |
| 3507 | JIE GENG | 6338 | QIAN HU | 371 | Platycodon grandiflorum | Angelica decursiva |
| 7047 | DA HUANG | 7847 | FANG FENG | 371 | Rheum officinale | Saposhnikovia divaricata |
| 5374 | DING XIANG | 6801 | GAN CAO | 371 | Syzygium aromaticum | Glycyrrhiza uralensis |
| 1010 | YUAN ZHI | 5421 | FU SHEN | 371 | Polygala tenuifolia | Poria |
| 2538 | DANG GUI | 5258 | CANG ZHU | 370 | Angelica sinensis | Atractylodes lancea |
| 7301 | BAI ZHU | 7520 | XIANG FU | 370 | Atractylodes macrocephala | Cyperus rotundus |
| 2522 | MAI DONG | 2538 | DANG GUI | 367 | Ophiopogon japonicus | Angelica sinensis |
| 5151 | MU XIANG | 7648 | HUANG LIAN | 365 | Saussurea lappa | Coptis chinensis |
| 3861 | REN SHEN | 5374 | DING XIANG | 364 | Panax ginseng | Syzygium aromaticum |
| 1849 | CHEN PI | 6919 | HUANG QI | 362 | Pericarpium Citri Reticulatae | Astragalus membranaceus |
| 1656 | GAN JIANG | 5800 | BAN XIA | 362 | Zingiber officinale | Pinellia ternata |
| 2538 | DANG GUI | 6079 | TAO REN | 358 | Angelica sinensis | Prunus persica |

### Supplementary Table 3. Description of the 268 literature-curated herb pairs.

| TCMID Herb_1 id | Herb_1 Pinyin name | TCMID Herb_2 id | Herb_2 Pinyin name | Herb_1 Latin name | Herb_2 Latin name |
|---|---|---|---|---|---|
| 1660 | MA HUANG | 3977 | GUI ZHI | Ephedra sinica | Cinnamomum cassia |
| 1932 | BING PIAN | 3811 | TIAN NAN XING | Dryobalanops aromatica | Arisaema consanguineum |
| 3861 | REN SHEN | 6919 | HUANG QI | Panax ginseng | Astragalus membranaceus |
| 3006 | SHENG MA | 3861 | REN SHEN | Cimicifuga foetida | Panax ginseng |
| 1509 | JUE MING ZI | 7307 | GOU QI ZI | Cassia tora | Lycium chinense |
| 1932 | BING PIAN | 4428 | SHE XIANG | Dryobalanops aromatica | Moschus moschiferus, Moschus berezovskii, Moschus sifanicus |
| 3861 | REN SHEN | 6604 | LIAN ZI | Panax ginseng | Nelumbo nucifera |
| 2522 | MAI DONG | 3861 | REN SHEN | Ophiopogon japonicus | Panax ginseng |
| 3861 | REN SHEN | 5894 | SHU DI HUANG | Panax ginseng | Rehmannia glutinosa |
| 3861 | REN SHEN | 5646 | GE JIE | Panax ginseng | Gecko |
| 6424 | ZHEN ZHU MU | 7282 | SHI JUE MING | Concha Margaritifera Usta;Margarita | Concha Haliotidis |
| 4287 | MU LI | 5310 | GUI BAN | Crassostrea gigas | Plastrum Testudinis |
| 1893 | BIE JIA | 4287 | MU LI | Carapax Trionycis | Crassostrea gigas |
| 1483 | CI SHI | 7282 | SHI JUE MING | Magnetitum | Concha Haliotidis |
| 1010 | YUAN ZHI | 6272 | SUAN ZAO REN | Polygala tenuifolia | Ziziphus jujuba var. spinosa |
| 6741 | TIAN MA | 7305 | GOU TENG | Gastrodia elata | Uncaria rhynchophylla |
| 1509 | JUE MING ZI | 2802 | BAI JI LI | Cassia tora | Fructus Tribuli |
| 3811 | TIAN NAN XING | 6741 | TIAN MA | Arisaema consanguineum | Gastrodia elata |
| 5800 | BAN XIA | 6741 | TIAN MA | Pinellia ternata | Gastrodia elata |
| 3977 | GUI ZHI | 6919 | HUANG QI | Cinnamomum cassia | Astragalus membranaceus |
| 1518 | WU WEI ZI | 6919 | HUANG QI | Schisandra chinensis | Astragalus membranaceus |
| 1345 | DANG SHEN | 6919 | HUANG QI | Codonopsis pilosula | Astragalus membranaceus |
| 2538 | DANG GUI | 6919 | HUANG QI | Angelica sinensis | Astragalus membranaceus |
| 6919 | HUANG QI | 7524 | ROU GUI | Astragalus membranaceus | Cinnamomum cassia |

| | | | | | |
|---|---|---|---|---|---|
| 3977 | GUI ZHI | 6801 | GAN CAO | Cinnamomum cassia | Glycyrrhiza uralensis |
| 3997 | BAI SHAO | 6801 | GAN CAO | Paeonia albiflora | Glycyrrhiza uralensis |
| 4063 | LU RONG | 5894 | SHU DI HUANG | Cervus nippon , Cervus elaphus | Rehmannia glutinosa |
| 1076 | DA ZAO | 6801 | GAN CAO | Ziziphus jujuba | Glycyrrhiza uralensis |
| 1656 | GAN JIANG | 3861 | REN SHEN | Zingiber officinale | Panax ginseng |
| 3861 | REN SHEN | 4063 | LU RONG | Panax ginseng | Cervus nippon , Cervus elaphus |
| 1345 | DANG SHEN | 7870 | FU LING | Codonopsis pilosula | Poria cocos |
| 2525 | ZI HE CHE | 3861 | REN SHEN | Homo sapiens | Panax ginseng |
| 6919 | HUANG QI | 7870 | FU LING | Astragalus membranaceus | Poria cocos |
| 1173 | FU ZI | 6919 | HUANG QI | Aconitum carmichaeli | Astragalus membranaceus |
| 3006 | SHENG MA | 6919 | HUANG QI | Cimicifuga foetida | Astragalus membranaceus |
| 6801 | GAN CAO | 6919 | HUANG QI | Glycyrrhiza uralensis | Astragalus membranaceus |
| 2538 | DANG GUI | 7382 | YI MU CAO | Angelica sinensis | Leonurus heterophyllus |
| 2474 | HONG HUA | 6079 | TAO REN | Carthamus tinctorius | Prunus persica |
| 6079 | TAO REN | 7047 | DA HUANG | Prunus persica | Rheum officinale |
| 5926 | CHUAN XIONG | 7520 | XIANG FU | Ligusticum chuanxiong | Cyperus rotundus |
| 2834 | RU XIANG | 4741 | MO YAO | Boswellia carterii | Commiphora myrrha |
| 1123 | ZE LAN | 2538 | DANG GUI | Lycopus lucidus | Angelica sinensis |
| 3796 | ZHE CHONG | 6636 | SHUI ZHI | Eupolyphaga seu steleophaga | Gardenia jasminoides var. grandiflora |
| 5800 | BAN XIA | 7484 | SHENG JIANG | Pinellia ternata | Zingiber officinale |
| 3303 | PU HUANG | 5354 | WU LING ZHI | Typha angustata | Trogopterus xanthipes, Pteromys volans |
| 2447 | BAI JI | 941 | E JIAO | Bletilla striata | Colla Corii Asini |
| 3303 | PU HUANG | 941 | E JIAO | Typha angustata | Colla Corii Asini |
| 3303 | PU HUANG | 7058 | QING DAI | Typha angustata | Indigo Naturalis |
| 4256 | SHENG DI HUANG | 4272 | CE BAI YE | Radix Rehmanniae | Thuja orientalis |
| 3997 | BAI SHAO | 4272 | CE BAI YE | Paeonia albiflora | Thuja orientalis |
| 3997 | BAI SHAO | 5926 | CHUAN XIONG | Paeonia albiflora | Ligusticum chuanxiong |

| | | | | | |
|---|---|---|---|---|---|
| 2538 | DANG GUI | 5926 | CHUAN XIONG | Angelica sinensis | Ligusticum chuanxiong |
| 1569 | XUE JIE | 2443 | SAN QI | Sanguis Draconis | Panax pseudo - ginseng var. notoginseng |
| 1483 | CI SHI | 5260 | ZHU SHA | Magnetitum | Cinnabaris |
| 5260 | ZHU SHA | 6123 | HU PO | Cinnabaris | Succinus |
| 4287 | MU LI | 6790 | LONG GU | Crassostrea gigas | Os Draconis |
| 1660 | MA HUANG | 7069 | BAI GUO | Ephedra sinica | Ginkgo biloba |
| 5260 | ZHU SHA | 7648 | HUANG LIAN | Cinnabaris | Coptis chinensis |
| 1010 | YUAN ZHI | 6161 | SHI CHANG PU | Polygala tenuifolia | Acorus tatarinowii |
| 2538 | DANG GUI | 6272 | SUAN ZAO REN | Angelica sinensis | Ziziphus jujuba var. spinosa |
| 1282 | BAI ZI REN | 6272 | SUAN ZAO REN | Biota orientalis | Ziziphus jujuba var. spinosa |
| 3811 | TIAN NAN XING | 5800 | BAN XIA | Arisaema consanguineum | Pinellia ternata |
| 5800 | BAN XIA | 7446 | XUAN FU HUA | Pinellia ternata | Flos Inulae |
| 1304 | ZHI MU | 6371 | CHUAN BEI MU | Anemarrhena asphodeloides | Fritillaria cirrhosa |
| 3861 | REN SHEN | 5800 | BAN XIA | Panax ginseng | Pinellia ternata |
| 5800 | BAN XIA | 6700 | HUANG QIN | Pinellia ternata | Scutellaria baicalensis |
| 1076 | DA ZAO | 4221 | TING LI ZI | Ziziphus jujuba | Lepidium apetalum |
| 1464 | ZI SU ZI | 7419 | XING REN | Fructus Perillae | Prunus armeniaca |
| 2074 | KUAN DONG HUA | 2215 | BAI HE | Tussilago farfara | Lilium brownii var. viridulum |
| 5310 | GUI BAN | 5736 | NIU XI | Plastrum Testudinis | Achyranthes bidentata |
| 1893 | BIE JIA | 5310 | GUI BAN | Carapax Trionycis | Plastrum Testudinis |
| 5310 | GUI BAN | 8114 | HUANG BAI | Plastrum Testudinis | Phellodendron amurense |
| 7307 | GOU QI ZI | 7682 | JU HUA | Lycium chinense | Chrysanthemum morifolium |
| 1033 | NV ZHEN ZI | 7307 | GOU QI ZI | Ligustrum lucidum | Lycium chinense |
| 3655 | LU JIAO JIAO | 8182 | GUI BAN JIAO | Pulvis Cornu Cervi | Colla carapacis et plastri testudinis |
| 7648 | HUANG LIAN | 7932 | HE ZI | Coptis chinensis | Terminalia chebula |
| 5279 | WU MEI | 6842 | YING SU KE | Prunus mume | Papaver somniferum |
| 1656 | GAN JIANG | 7932 | HE ZI | Zingiber officinale | Terminalia chebula |
| 1518 | WU WEI ZI | 6584 | WU BEI ZI | Schisandra chinensis | Galla Chinensis |

| | | | | | |
|---|---|---|---|---|---|
| 1369 | TIAN DONG | 4256 | SHENG DI HUANG | Radix Asparagi | Radix Rehmanniae |
| 1369 | TIAN DONG | 5894 | SHU DI HUANG | Radix Asparagi | Rehmannia glutinosa |
| 2522 | MAI DONG | 8088 | YU ZHU | Ophiopogon japonicus | Polygonatum odoratum |
| 2522 | MAI DONG | 4256 | SHENG DI HUANG | Ophiopogon japonicus | Radix Rehmanniae |
| 2522 | MAI DONG | 5034 | XUAN SHEN | Ophiopogon japonicus | Scrophularia ningpoensis |
| 3861 | REN SHEN | 7307 | GOU QI ZI | Panax ginseng | Lycium chinense |
| 5404 | TU SI ZI | 7307 | GOU QI ZI | Semen Cuseutae;Semen Cuseutae | Lycium chinense |
| 1304 | ZHI MU | 2215 | BAI HE | Anemarrhena asphodeloides | Lilium brownii var. viridulum |
| 3841 | GUA DI | 5082 | CHI XIAO DOU | Cucumis melo | Semen Phaseoli |
| 6238 | XIONG HUANG | 7399 | SHE CHUANG ZI | Realgar | Cnidium monnieri |
| 1932 | BING PIAN | 5196 | LU GAN SHI | Dryobalanops aromatica | Calamina |
| 1387 | SANG PIAO XIAO | 5009 | FU PEN ZI | Ootheca Mantidis | Rubus idaeus |
| 1702 | SHA YUAN ZI | 5009 | FU PEN ZI | Semen Astragali Complanati | Rubus idaeus |
| 7399 | SHE CHUANG ZI | 7741 | KU SHEN | Cnidium monnieri | Sophora flavescens |
| 2033 | MANG XIAO | 5260 | ZHU SHA | Natrii Sulfas1 | Cinnabaris |
| 2033 | MANG XIAO | 3192 | PENG SHA | Natrii Sulfas1 | Sal Sedatirum |
| 1932 | BING PIAN | 3192 | PENG SHA | Dryobalanops aromatica | Sal Sedatirum |
| 1656 | GAN JIANG | 6358 | CHI SHI ZHI | Zingiber officinale | Halloysitum Rubrum |
| 6510 | QIAN SHI | 6604 | LIAN ZI | Semen Euryales | Nelumbo nucifera |
| 1630 | JIN YING ZI | 6510 | QIAN SHI | Rosa laevigata | Semen Euryales |
| 5784 | YU YU LIANG | 6358 | CHI SHI ZHI | Limonitum | Halloysitum Rubrum |
| 1387 | SANG PIAO XIAO | 3861 | REN SHEN | Ootheca Mantidis | Panax ginseng |
| 1387 | SANG PIAO XIAO | 6790 | LONG GU | Ootheca Mantidis | Os Draconis |
| 1630 | JIN YING ZI | 5009 | FU PEN ZI | Rosa laevigata | Rubus idaeus |
| 2525 | ZI HE CHE | 5894 | SHU DI HUANG | Homo sapiens | Rehmannia glutinosa |

| | | | | | |
|---|---|---|---|---|---|
| 1702 | SHA YUAN ZI | 5404 | TU SI ZI | Semen Astragali Complanati | Semen Cuseutae;Semen Cuseutae |
| 2538 | DANG GUI | 5894 | SHU DI HUANG | Angelica sinensis | Rehmannia glutinosa |
| 5948 | BU GU ZHI | 6842 | YING SU KE | Psoralea corylifolia | Papaver somniferum |
| 1387 | SANG PIAO XIAO | 7655 | YI ZHI REN | Ootheca Mantidis | Alpinia oxyphylla |
| 2525 | ZI HE CHE | 5310 | GUI BAN | Homo sapiens | Plastrum Testudinis |
| 5310 | GUI BAN | 5894 | SHU DI HUANG | Plastrum Testudinis | Rehmannia glutinosa |
| 2538 | DANG GUI | 941 | E JIAO | Angelica sinensis | Colla Corii Asini |
| 5894 | SHU DI HUANG | 6859 | SHAN ZHU YU | Rehmannia glutinosa | Cornus officinalis |
| 1173 | FU ZI | 2538 | DANG GUI | Aconitum carmichaeli | Angelica sinensis |
| 2538 | DANG GUI | 4063 | LU RONG | Angelica sinensis | Cervus nippon , Cervus elaphus |
| 1173 | FU ZI | 4063 | LU RONG | Aconitum carmichaeli | Cervus nippon , Cervus elaphus |
| 1883 | ROU CONG RONG | 3014 | SUO YANG | Cistanche deserticola | Cynomorium songaricum |
| 4063 | LU RONG | 6859 | SHAN ZHU YU | Cervus nippon , Cervus elaphus | Cornus officinalis |
| 5948 | BU GU ZHI | 7482 | HU TAO REN | Psoralea corylifolia | Juglans regia |
| 2982 | YIN YANG HUO | 7589 | XIAN MAO | Epimedium brevicornum | Curculigo orchioides |
| 6802 | BA JI TIAN | 7589 | XIAN MAO | Morinda officinalis | Curculigo orchioides |
| 3997 | BAI SHAO | 5310 | GUI BAN | Paeonia albiflora | Plastrum Testudinis |
| 6919 | HUANG QI | 941 | E JIAO | Astragalus membranaceus | Colla Corii Asini |
| 3861 | REN SHEN | 941 | E JIAO | Panax ginseng | Colla Corii Asini |
| 2538 | DANG GUI | 3997 | BAI SHAO | Angelica sinensis | Paeonia albiflora |
| 3997 | BAI SHAO | 7307 | GOU QI ZI | Paeonia albiflora | Lycium chinense |
| 5954 | LONG YAN ROU | 6272 | SUAN ZAO REN | Arillus Longan | Ziziphus jujuba var. spinosa |
| 1518 | WU WEI ZI | 2522 | MAI DONG | Schisandra chinensis | Ophiopogon japonicus |
| 2522 | MAI DONG | 5800 | BAN XIA | Ophiopogon japonicus | Pinellia ternata |
| 1369 | TIAN DONG | 2522 | MAI DONG | Radix Asparagi | Ophiopogon japonicus |

| | | | | | |
|---|---|---|---|---|---|
| 3997 | BAI SHAO | 5894 | SHU DI HUANG | Paeonia albiflora | Rehmannia glutinosa |
| 3861 | REN SHEN | 6281 | HE SHOU WU | Panax ginseng | Polygonum multiflorum |
| 5894 | SHU DI HUANG | 6281 | HE SHOU WU | Rehmannia glutinosa | Polygonum multiflorum |
| 1304 | ZHI MU | 5894 | SHU DI HUANG | Anemarrhena asphodeloides | Rehmannia glutinosa |
| 3997 | BAI SHAO | 4287 | MU LI | Paeonia albiflora | Crassostrea gigas |
| 2802 | BAI JI LI | 6281 | HE SHOU WU | Fructus Tribuli | Polygonum multiflorum |
| 2538 | DANG GUI | 6281 | HE SHOU WU | Angelica sinensis | Polygonum multiflorum |
| 2221 | ZHI ZI | 5512 | HUAI HUA | Gardenia jasminoides | flos Sophorae |
| 7648 | HUANG LIAN | 7935 | WU ZHU YU | Coptis chinensis | Evodia rutaecarpa |
| 6205 | GE GEN | 7648 | HUANG LIAN | Pueraria lobata | Coptis chinensis |
| 7047 | DA HUANG | 7648 | HUANG LIAN | Rheum officinale | Coptis chinensis |
| 2221 | ZHI ZI | 6700 | HUANG QIN | Gardenia jasminoides | Scutellaria baicalensis |
| 2522 | MAI DONG | 7648 | HUANG LIAN | Ophiopogon japonicus | Coptis chinensis |
| 5279 | WU MEI | 7648 | HUANG LIAN | Prunus mume | Coptis chinensis |
| 3379 | ZHI SHI | 7648 | HUANG LIAN | Citrus aurantium | Coptis chinensis |
| 2057 | PU GONG YING | 2475 | JIN YIN HUA | Taraxacum mongolicum | Lonicera japonica |
| 5151 | MU XIANG | 7648 | HUANG LIAN | Saussurea lappa | Coptis chinensis |
| 1304 | ZHI MU | 8114 | HUANG BAI | Anemarrhena asphodeloides | Phellodendron amurense |
| 7648 | HUANG LIAN | 8114 | HUANG BAI | Coptis chinensis | Phellodendron amurense |
| 5258 | CANG ZHU | 8114 | HUANG BAI | Atractylodes lancea | Phellodendron amurense |
| 1656 | GAN JIANG | 7648 | HUANG LIAN | Zingiber officinale | Coptis chinensis |
| 3861 | REN SHEN | 7648 | HUANG LIAN | Panax ginseng | Coptis chinensis |
| 2221 | ZHI ZI | 8114 | HUANG BAI | Gardenia jasminoides | Phellodendron amurense |
| 1304 | ZHI MU | 6700 | HUANG QIN | Anemarrhena asphodeloides | Scutellaria baicalensis |
| 6700 | HUANG QIN | 7648 | HUANG LIAN | Scutellaria baicalensis | Coptis chinensis |

| | | | | | |
|---|---|---|---|---|---|
| 6999 | BAI TOU WENG | 8114 | HUANG BAI | Pulsatilla chinensis | Phellodendron amurense |
| 1957 | CHI SHAO | 3997 | BAI SHAO | Radix Paeoniae Rubra | Paeonia albiflora |
| 1957 | CHI SHAO | 5926 | CHUAN XIONG | Radix Paeoniae Rubra | Ligusticum chuanxiong |
| 4256 | SHENG DI HUANG | 5034 | XUAN SHEN | Radix Rehmanniae | Scrophularia ningpoensis |
| 1419 | SHAN DOU GEN | 5207 | SHE GAN | Sophora subprostrata | Belamcanda chinensis |
| 6196 | QING PI | 6999 | BAI TOU WENG | Pericarpium Citri Reticulatae Viride | Pulsatilla chinensis |
| 3006 | SHENG MA | 4256 | SHENG DI HUANG | Cimicifuga foetida | Radix Rehmanniae |
| 4256 | SHENG DI HUANG | 5894 | SHU DI HUANG | Radix Rehmanniae | Rehmannia glutinosa |
| 3006 | SHENG MA | 5034 | XUAN SHEN | Cimicifuga foetida | Scrophularia ningpoensis |
| 3997 | BAI SHAO | 4256 | SHENG DI HUANG | Paeonia albiflora | Radix Rehmanniae |
| 2057 | PU GONG YING | 8202 | XIA KU CAO | Taraxacum mongolicum | Prunella vulgaris |
| 2057 | PU GONG YING | 6597 | ZI HUA DI DING | Taraxacum mongolicum | Viola yedoensis |
| 1050 | YE JU HUA | 2057 | PU GONG YING | Chrysanthemum indicum | Taraxacum mongolicum |
| 3673 | NIU BANG ZI | 4518 | LIAN QIAO | Arctium lappa | Forsythia suspensa |
| 1932 | BING PIAN | 7058 | QING DAI | Dryobalanops aromatica | Indigo Naturalis |
| 2756 | TU FU LING | 3445 | BAI XIAN PI | Smilax glabra | Cortex dictamni |
| 5107 | SHI GAO | 7869 | DA QING YE | Gypsum Fibrosum | Isatis indigotica |
| 3370 | BAN LAN GEN | 5034 | XUAN SHEN | Isatis indigotica | Scrophularia ningpoensis |
| 4776 | QIANG HUO | 7847 | FANG FENG | Notopterygium incisum | Saposhnikovia divaricata |
| 4776 | QIANG HUO | 5926 | CHUAN XIONG | Notopterygium incisum | Ligusticum chuanxiong |
| 4449 | DU HUO | 4776 | QIANG HUO | Angelica pubescens f. biserrata | Notopterygium incisum |
| 4526 | JING JIE | 7847 | FANG FENG | Schizonepeta tenuifolia | Saposhnikovia divaricata |
| 6919 | HUANG QI | 7847 | FANG FENG | Astragalus membranaceus | Saposhnikovia divaricata |

| | | | | | |
|---|---|---|---|---|---|
| 1428 | CONG BAI | 7484 | SHENG JIANG | Allium fistulosum | Zingiber officinale |
| 1076 | DA ZAO | 7484 | SHENG JIANG | Ziziphus jujuba | Zingiber officinale |
| 3977 | GUI ZHI | 6205 | GE GEN | Cinnamomum cassia | Pueraria lobata |
| 1602 | XIN YI | 6163 | CANG ER ZI | Magnolia liliflora | Fructus Xanthii sibirici |
| 1660 | MA HUANG | 7484 | SHENG JIANG | Ephedra sinica | Zingiber officinale |
| 1660 | MA HUANG | 5107 | SHI GAO | Ephedra sinica | Gypsum Fibrosum |
| 1660 | MA HUANG | 6205 | GE GEN | Ephedra sinica | Pueraria lobata |
| 1660 | MA HUANG | 7419 | XING REN | Ephedra sinica | Prunus armeniaca |
| 3977 | GUI ZHI | 5926 | CHUAN XIONG | Cinnamomum cassia | Ligusticum chuanxiong |
| 3977 | GUI ZHI | 3997 | BAI SHAO | Cinnamomum cassia | Paeonia albiflora |
| 1660 | MA HUANG | 6801 | GAN CAO | Ephedra sinica | Glycyrrhiza uralensis |
| 3861 | REN SHEN | 5107 | SHI GAO | Panax ginseng | Gypsum Fibrosum |
| 5107 | SHI GAO | 5258 | CANG ZHU | Gypsum Fibrosum | Atractylodes lancea |
| 5107 | SHI GAO | 5894 | SHU DI HUANG | Gypsum Fibrosum | Rehmannia glutinosa |
| 3673 | NIU BANG ZI | 3768 | FU PING | Arctium lappa | Lemna minor |
| 1304 | ZHI MU | 5107 | SHI GAO | Anemarrhena asphodeloides | Gypsum Fibrosum |
| 7524 | ROU GUI | 7648 | HUANG LIAN | Cinnamomum cassia | Coptis chinensis |
| 2221 | ZHI ZI | 5790 | GAO LIANG JIANG | Gardenia jasminoides | Alpinia officinarum |
| 2150 | DAN DOU CHI | 2221 | ZHI ZI | Semen Sojae Praeparata | Gardenia jasminoides |
| 2356 | SANG YE | 5020 | SANG ZHI | Morus alba | Morus alba |
| 5926 | CHUAN XIONG | 7682 | JU HUA | Ligusticum chuanxiong | Chrysanthemum morifolium |
| 7682 | JU HUA | 7847 | FANG FENG | Chrysanthemum morifolium | Saposhnikovia divaricata |
| 2356 | SANG YE | 7682 | JU HUA | Morus alba | Chrysanthemum morifolium |
| 3006 | SHENG MA | 6205 | GE GEN | Cimicifuga foetida | Pueraria lobata |
| 2802 | BAI JI LI | 7428 | MAN JING ZI | Fructus Tribuli | Vitex trifolia |
| 7428 | MAN JING ZI | 7682 | JU HUA | Vitex trifolia | Chrysanthemum morifolium |
| 2475 | JIN YIN HUA | 4518 | LIAN QIAO | Lonicera japonica | Forsythia suspensa |
| 1849 | CHEN PI | 5800 | BAN XIA | Pericarpium Citri Reticulatae | Pinellia ternata |

| | | | | | |
|---|---|---|---|---|---|
| 3379 | ZHI SHI | 5050 | HOU PU | Citrus aurantium | Cortex Magnoliae officinalis |
| 2896 | BING LANG | 5151 | MU XIANG | Areca catechu | Saussurea lappa |
| 1098 | HU LU BA | 6331 | XIAO HUI XIANG | Trigonella foenum - graecum | fructus Foeniculi |
| 1767 | CAO WU | 3194 | CHUAN WU | Radix Aconiti Kusnezoffii | Radix Aconiti |
| 2538 | DANG GUI | 7520 | XIANG FU | Angelica sinensis | Cyperus rotundus |
| 1397 | ZI SU GENG | 7520 | XIANG FU | Perilla frutescens var. arguta | Cyperus rotundus |
| 5790 | GAO LIANG JIANG | 7520 | XIANG FU | Alpinia officinarum | Cyperus rotundus |
| 5151 | MU XIANG | 7047 | DA HUANG | Saussurea lappa | Rheum officinale |
| 5151 | MU XIANG | 6681 | WU YAO | Saussurea lappa | Lindera strychnifolia |
| 6331 | XIAO HUI XIANG | 7935 | WU ZHU YU | fructus Foeniculi | Evodia rutaecarpa |
| 2910 | SHI DI | 5374 | DING XIANG | Diospyros kaki | Syzygium aromaticum |
| 1656 | GAN JIANG | 5790 | GAO LIANG JIANG | Zingiber officinale | Alpinia officinarum |
| 2538 | DANG GUI | 7524 | ROU GUI | Angelica sinensis | Cinnamomum cassia |
| 1656 | GAN JIANG | 7524 | ROU GUI | Zingiber officinale | Cinnamomum cassia |
| 3379 | ZHI SHI | 6621 | XIE BAI | Citrus aurantium | Allium macrostemon |
| 2202 | SHI JUN ZI | 3683 | LU HUI | Quisqualis indica | Aloe vera |
| 2896 | BING LANG | 4848 | QIAN NIU ZI | Areca catechu | Pharbitis nil |
| 2502 | CHUAN LIAN ZI | 3993 | YAN HU SUO | Melia toosendan | Corydalis yanhusuo |
| 2612 | GUA LOU | 6621 | XIE BAI | Trichosanthes kirilowii | Allium macrostemon |
| 4256 | SHENG DI HUANG | 6014 | BAI MAO GEN | Radix Rehmanniae | Rhizoma Imperatae |
| 5512 | HUAI HUA | 6800 | DI YU | flos Sophorae | Sanguisorba officinalis |
| 2697 | NAN GUA ZI | 2896 | BING LANG | Cucurbita moschata | Areca catechu |
| 5151 | MU XIANG | 7520 | XIANG FU | Saussurea lappa | Cyperus rotundus |
| 6681 | WU YAO | 7655 | YI ZHI REN | Lindera strychnifolia | Alpinia oxyphylla |
| 6681 | WU YAO | 6726 | CHEN XIANG | Lindera strychnifolia | Aquilaria agallocha |
| 3997 | BAI SHAO | 7520 | XIANG FU | Paeonia albiflora | Cyperus rotundus |
| 5151 | MU XIANG | 6726 | CHEN XIANG | Saussurea lappa | Aquilaria agallocha |
| 5374 | DING XIANG | 6726 | CHEN XIANG | Syzygium aromaticum | Aquilaria agallocha |
| 6681 | WU YAO | 7520 | XIANG FU | Lindera strychnifolia | Cyperus rotundus |

| | | | | | |
|---|---|---|---|---|---|
| 3336 | GAN SUI | 7504 | YUAN HUA | Euphorbia kansui | Daphne genkwa |
| 4848 | QIAN NIU ZI | 7047 | DA HUANG | Pharbitis nil | Rheum officinale |
| 4449 | DU HUO | 6690 | SANG JI SHENG | Angelica pubescens f. biserrata | Loranthus parasiticus |
| 5050 | HOU PU | 7047 | DA HUANG | Cortex Magnoliae officinalis | Rheum officinale |
| 6070 | HUO MA REN | 7422 | YU LI REN | Cannabis sativa | Prunus japonica |
| 6079 | TAO REN | 7419 | XING REN | Prunus persica | Prunus armeniaca |
| 5050 | HOU PU | 5258 | CANG ZHU | Cortex Magnoliae officinalis | Atractylodes lancea |
| 2338 | BAI HUA SHE | 5145 | WU SHAO SHE | Bungarus Parvus | Zaocys |
| 1849 | CHEN PI | 5258 | CANG ZHU | Pericarpium Citri Reticulatae | Atractylodes lancea |
| 4696 | FANG JI | 6919 | HUANG QI | Stephania tetrandra | Astragalus membranaceus |
| 6700 | HUANG QIN | 7616 | QING HAO | Scutellaria baicalensis | Artemisia apiacea |
| 1304 | ZHI MU | 5106 | DI GU PI | Anemarrhena asphodeloides | Cortex Lycii Radicis |
| 2696 | YIN CHAI HU | 3706 | HU HUANG LIAN | Stellaria dichotoma var. lanceolata | Picrorhiza kurrooa |
| 1893 | BIE JIA | 7616 | QING HAO | Carapax Trionycis | Artemisia apiacea |
| 1173 | FU ZI | 7047 | DA HUANG | Aconitum carmichaeli | Rheum officinale |
| 3379 | ZHI SHI | 7047 | DA HUANG | Citrus aurantium | Rheum officinale |
| 2033 | MANG XIAO | 7047 | DA HUANG | Natrii Sulfas1 | Rheum officinale |
| 1173 | FU ZI | 3977 | GUI ZHI | Aconitum carmichaeli | Cinnamomum cassia |
| 1173 | FU ZI | 1656 | GAN JIANG | Aconitum carmichaeli | Zingiber officinale |
| 1173 | FU ZI | 3861 | REN SHEN | Aconitum carmichaeli | Panax ginseng |
| 1173 | FU ZI | 7524 | ROU GUI | Aconitum carmichaeli | Cinnamomum cassia |
| 2683 | QU MAI | 7426 | BIAN XU | Dianthus superbus | Polygonum aviculare |
| 1173 | FU ZI | 5894 | SHU DI HUANG | Aconitum carmichaeli | Rehmannia glutinosa |
| 1656 | GAN JIANG | 5800 | BAN XIA | Zingiber officinale | Pinellia ternata |
| 1173 | FU ZI | 6801 | GAN CAO | Aconitum carmichaeli | Glycyrrhiza uralensis |

| | | | | | |
|---|---|---|---|---|---|
| 1173 | FU ZI | 3997 | BAI SHAO | Aconitum carmichaeli | Paeonia albiflora |
| 1472 | WANG BU LIU XING | 2276 | MU TONG | Vaccaria segetalis | Akebia quinata |
| 4660 | HUA SHI | 6801 | GAN CAO | Talcum | Glycyrrhiza uralensis |
| 4883 | ZHU LING | 7870 | FU LING | Polyporus umbellatus | Poria cocos |
| 3977 | GUI ZHI | 7870 | FU LING | Cinnamomum cassia | Poria cocos |
| 4981 | BI XIE | 7655 | YI ZHI REN | Dioscorea hypoglauca | Alpinia oxyphylla |
| 4783 | HAI JIN SHA | 7734 | SHI WEI | Lygodium japonicum | Pyrrosia lingua |
| 4783 | HAI JIN SHA | 7238 | JIN QIAN CAO | Lygodium japonicum | Herba Glechomae Longitubae |

### Supplementary Table 4. Performance of the network models on the 268 literature-curated herb pairs.

| Ingredient-level distance type | Target-level distance type | Distance for random herb pairs | Distance for top herb pairs | p-value | AUROC | AUPRC |
|---|---|---|---|---|---|---|
| center | center | 2.0478542 | 1.74093754 | 2.11E-09 | 0.65258255 | 0.66995736 |
| center | closest | 2.13326842 | 1.75781903 | 3.36E-14 | 0.72778048 | 0.70564243 |
| center | kernel | 3.35811247 | 3.04950001 | 2.08E-12 | 0.73497053 | 0.71112342 |
| center | separation | 0.47527221 | -0.0750626 | 1.01E-14 | 0.73184382 | 0.69605913 |
| center | shortest | 2.40570581 | 2.06997944 | 1.35E-14 | 0.75941309 | 0.73781801 |
| closest | center | 2.05668841 | 1.73543657 | 9.68E-16 | 0.75145514 | 0.72128538 |
| closest | closest | 2.15539272 | 1.83480915 | 1.23E-17 | 0.75261301 | 0.74184792 |
| closest | kernel | 3.40006316 | 3.17832449 | 8.32E-13 | 0.71297054 | 0.72377225 |
| closest | separation | 0.55887054 | 0.1363911 | 2.80E-12 | 0.71282406 | 0.69255095 |
| closest | shortest | 2.46641736 | 2.26684315 | 7.59E-11 | 0.68774285 | 0.71234357 |
| kernel | center | 3.13641665 | 3.06609757 | 0.05468342 | 0.55303022 | 0.59022393 |
| kernel | closest | 3.24234786 | 3.18873124 | 0.10054827 | 0.54253952 | 0.60750301 |
| kernel | kernel | 4.47407609 | 4.44833525 | 0.35181911 | 0.50115185 | 0.59125441 |
| kernel | separation | 1.67543428 | 1.56901708 | 0.07307494 | 0.51081052 | 0.58346722 |
| kernel | shortest | 3.54028704 | 3.51964589 | 0.40551305 | 0.49688732 | 0.5905264 |
| separation | center | 1.12212451 | 0.50368471 | 4.48E-16 | 0.71391659 | 0.74067945 |
| separation | closest | 1.2184745 | 0.62582169 | 2.99E-15 | 0.70613178 | 0.74215356 |
| separation | kernel | 1.69715839 | 0.55630479 | 2.02E-27 | 0.73413851 | 0.77375432 |
| separation | separation | 0.51090354 | 0.45942857 | 0.37262747 | 0.48424361 | 0.54172542 |
| separation | shortest | 1.23828789 | 0.38080579 | 1.37E-26 | 0.73570036 | 0.77593863 |
| shortest | center | 2.15526282 | 2.12164611 | 0.2995525 | 0.49200057 | 0.55605116 |
| shortest | closest | 2.25776206 | 2.24052633 | 0.46051436 | 0.49847783 | 0.57431898 |
| shortest | kernel | 3.48478307 | 3.47814613 | 0.53018203 | 0.48086922 | 0.57374846 |
| shortest | separation | 0.71103195 | 0.67811359 | 0.46434602 | 0.47292169 | 0.55393498 |
| shortest | shortest | 2.55074643 | 2.54703297 | 0.51184335 | 0.48047365 | 0.57613159 |

### Supplementary Table 5. Description of *Astragalus membranaceus* and *Glycyrrhiza uralensis*.

| TCMID herb pinyin name | TCMID herb latine name | TCMID herb id | TCMID herb Website | TCMID ingredient_name | TCMID ingredient id | TCMID Ingredient Website | PubChem id | PubChem ID website | Pubchem_inchikey | Pubchem_molecular_formula | Pubchem_smiles | Stitch_cid_m | Stitch_cid_s | Stitch name |
|---|---|---|---|---|---|---|---|---|---|---|---|---|---|---|
| GANCAO | Glycyrrhiza uralensis | 6801 | http://119.3.41.228:8000/tcmid/herb/6801/ | ganoderic acid a | 27722 | http://119.3.41.228:8000/tcmid/ingredient/27722/ | 14193987 | http://pubchem.ncbi.nlm.nih.gov/compound/14193987 | OVUOUFPIPZJGME-JQDIJSAQSA-N | C30H42O7 | CC(CC(=O)C=C(C)C1CC(C2(C1(CC(=O)C3=C2CC(CC4C3(CCC(=O)C4(C)C)O)C)C)O)C)C(=O)O | CIDm64420888 | CIDs14193987 | AC1O5WK1 |
| GANCAO | Glycyrrhiza uralensis | 6801 | http://119.3.41.228:8000/tcmid/herb/6801/ | glycyrrhetinicacid | 8840 | http://119.3.41.228:8000/tcmid/ingredient/8840/ | 5311038 | http://pubchem.ncbi.nlm.nih.gov/compound/5311038 | OBZHEBDUNPOCJG-ILYNEHRHSA-N | C34H50O7 | CC1(C2CCC3(C(C2(CCC1OC(=O)CCC(=O)O)C)C(=O)C=C4C3(CCC5(C4CC(CC5)(C)C(=O)O)C)C)C)C | CIDm00025561 | CIDs05311038 | Bio-Plex |
| GANCAO | Glycyrrhiza uralensis | 6801 | http://119.3.41.228:8000/tcmid/herb/6801/ | gamma-sitosterol | 29509 | http://119.3.41.228:8000/tcmid/ingredient/29509/ | 636741 | http://pubchem.ncbi.nlm.nih.gov/compound/636741 | KZJWDPNRJALLNS-JQEGLAGPSA-N | C29H50O | CCC(CCC(C)C1CCC2C1(CCC3C2CC=C4C3(CCC(C4)O)C)C)C(C)C | CIDm00006744 | CIDs00636741 | clionasterol |
| GANCAO | Glycyrrhiza uralensis | 6801 | http://119.3.41.228:8000/tcmid/herb/6801/ | β-sitosterol | 19967 | http://119.3.41.228:8000/tcmid/ingredient/19967/ | 222284 | http://pubchem.ncbi.nlm.nih.gov/compound/222284 | KZJWDPNRJALLNS-VJSFXXLFSA-N | C29H50O | CCC(CCC(C)C1CCC2C1(CCC3C2CC=C4C3(CCC(C4)O)C)C)C(C)C | CIDm00006744 | CIDs00222284 | clionasterol |
| GANCAO | Glycyrrhiza uralensis | 6801 | http://119.3.41.228:8000/tcmid/herb/6801/ | 2,5-dihydroxymethyl-3,4-dihydroxypyrrolidine | 6027 | http://119.3.41.228:8000/tcmid/ingredient/6027/ | 124702 | http://pubchem.ncbi.nlm.nih.gov/compound/124702 | PFYHYHZGDNWFIF-KVTDHHQDSA-N | C6H13NO4 | C(C1C(C(C(N1)CO)O)O)O | CIDm00001472 | CIDs00124702 | DB02172 |
| GANCAO | Glycyrrhiza uralensis | 6801 | http://119.3.41.228:8000/tcmid/herb/6801/ | ferulic acid | 23696 | http://119.3.41.228:8000/tcmid/ingredient/23696/ | 54691413 | http://pubchem.ncbi.nlm.nih.gov/compound/54691413 | KSEBMYQBYZTDHS-HWKANZROSA-M | C10H9O4- | COC1=C(C=CC(=C1)C=CC(=O)O)[O-] | CIDm00000709 | CIDs00445858 | ferulic acid |
| GANCAO | Glycyrrhiza urale | 6801 | http://119.3.41.228:8000/tcmid/herb/6801/ | formononetin | 7882 | http://119.3.41.228:8000/tcmid/ingredient/7882/ | 10378473 | http://pubchem.ncbi.nlm.nih.gov/compound/10378473 | HKQYGTCOTHHOMP-OPOMH | C16H12O4 | COC1=CC=C(C=C1)C2=COC3=C(C2=O)C=CC(=C3)O | CIDm52803788 | CIDs10378473 | formononetin |

| | | | | | | | | | | | | | | |
|---|---|---|---|---|---|---|---|---|---|---|---|---|---|---|
| | nsis | | | | | | 73 | | RLSSA-N | | | | | |
| GANCAO | Glycyrrhiza uralensis | 6801 | http://119.3.41.228:8000/tcmid/herb/6801/ | 18alpha-glycyrrhetinic acid | 23166 | http://119.3.41.228:8000/tcmid/ingredient/23166/ | 16219451 | http://pubchem.ncbi.nlm.nih.gov/compound/16219451 | MPDGHEJMBKOTSU-WWCMNKNLSA-N | C30H46O4 | CC1(C2CCC3(C(C2(CCC1O)C)C(=O)C=C4C3(CCC5(C4CC(CC5)(C)C(=O)O)C)C)C)C | CIDm00003230 | CIDs16219451 | glycyrrhetinic. |
| GANCAO | Glycyrrhiza uralensis | 6801 | http://119.3.41.228:8000/tcmid/herb/6801/ | 18beta-glycyrrhetinic acid | 23092 | http://119.3.41.228:8000/tcmid/ingredient/23092/ | 10114 | http://pubchem.ncbi.nlm.nih.gov/compound/10114 | MPDGHEJMBKOTSU-YKLVYJNSSA-N | C30H46O4 | CC1(C2CCC3(C(C2(CCC1O)C)C(=O)C=C4C3(CCC5(C4CC(CC5)(C)C(=O)O)C)C)C)C | CIDm00003230 | CIDs00010114 | glycyrrhetinic. |
| GANCAO | Glycyrrhiza uralensis | 6801 | http://119.3.41.228:8000/tcmid/herb/6801/ | glycyrrhetinic acid | 23291 | http://119.3.41.228:8000/tcmid/ingredient/23291/ | 18526330 | http://pubchem.ncbi.nlm.nih.gov/compound/18526330 | MPDGHEJMBKOTSU-WFJWTYAKSA-N | C30H46O4 | CC1(C2CCC3(C(C2(CCC1O)C)C(=O)C=C4C3(CCC5(C4CC(CC5)(C)C(=O)O)C)C)C)C | CIDm00003230 | CIDs18526330 | glycyrrhetinic. |
| GANCAO | Glycyrrhiza uralensis | 6801 | http://119.3.41.228:8000/tcmid/herb/6801/ | glycyrrhizic acid | 23251 | http://119.3.41.228:8000/tcmid/ingredient/23251/ | 46878350 | http://pubchem.ncbi.nlm.nih.gov/compound/46878350 | LPLVUJXQOOQHMX-YFINQFOSA-N | C42H62O16 | CC1(C2CCC3(C(C2(CCC1OC4C(C(C(C(O4)C(=O)O)O)O)OC5C(C(C(C(O5)C(=O)O)O)O)O)C)C(=O)C=C6C3(CCC7(C6CC(CC7)(C)C(=O)O)C)C)C)C | CIDm00003495 | CIDs46878350 | glycyrrhizin |
| GANCAO | Glycyrrhiza uralensis | 6801 | http://119.3.41.228:8000/tcmid/herb/6801/ | glycyrrhizic acid | 8845 | http://119.3.41.228:8000/tcmid/ingredient/8845/ | 128229 | http://pubchem.ncbi.nlm.nih.gov/compound/128229 | LPLVUJXQOOQHMX-MOGLOQIBSA-N | C42H62O16 | CC1(C2CCC3(C(C2(CCC1OC4C(C(C(C(O4)C(=O)O)O)O)OC5C(C(C(C(O5)C(=O)O)O)O)O)C)C(=O)C=C6C3(CCC7(C6CC(CC7)(C)C(=O)O)C)C)C)C | CIDm00003495 | CIDs00128229 | glycyrrhizin |
| GANCAO | Glycyrrhiza uralensis | 6801 | http://119.3.41.228:8000/tcmid/herb/6801/ | glycyrrhizin | 23172 | http://119.3.41.228:8000/tcmid/ingredient/23172/ | 46902095 | http://pubchem.ncbi.nlm.nih.gov/compound/46902095 | ILRKKHJEINIICQ-UIAZACTHSA-O | C42H66NO16+ | CC1(C2CCC3(C(C2(CCC1OC4C(C(C(C(O4)C(=O)O)O)O)OC5C(C(C(C(O5)C(=O)O)O)O)O)C)C(=O)C=C6C3(CCC7(C6CC(CC7)(C)C(=O)O)C)C)C)C.[NH4+] | CIDm00003495 | CIDs16213697 | glycyrrhizin |
| GANCAO | Glycyrrhiza uralensis | 6801 | http://119.3.41.228:8000/tcmid/herb/6801/ | monoammonium glycyrrhizinate | 23259 | http://119.3.41.228:8000/tcmid/ingredient/23259/ | 45357226 | http://pubchem.ncbi.nlm.nih.gov/compound/45357226 | ILRKKHJEINIICQ-UFTZEXNXSA-N | C42H65NO16 | CC1(C2CCC3(C(C2(CCC1OC4C(C(C(C(O4)C(=O)O)O)O)OC5C(C(C(C(O5)C(=O)O)O)O)O)C)C(=O)C=C6C3(CCC7(C6CC(CC7)(C)C(=O)O)C)C)C)C.N | CIDm00003495 | CIDs00451751 | glycyrrhizin |
| GANCAO | Glycyrrhiza uralensis | 6801 | http://119.3.41.228:8000/tcmid/herb/6801/ | hispidulin | 9563 | http://119.3.41.228:8000/tcmid/ingredient/9563/ | 5281628 | http://pubchem.ncbi.nlm.nih.gov/compound/5281628 | IHFBPDAQLQOCBX-UHFFFAOYSA-N | C16H12O6 | COC1=C(C=C2C(=C1O)C(=O)C=C(O2)C3=CC=C(C=C3)O)O | CIDm05281628 | CIDs05281628 | hispidulin |

| | | | | | | | | | | | | | |
|---|---|---|---|---|---|---|---|---|---|---|---|---|---|
| GANCAO | Glycyrrhiza uralensis | 6801 | http://119.3.41.228:8000/tcmid/herb/6801/ | isolicoflavonol | 11487 | http://119.3.41.228:8000/tcmid/ingredient/11487/ | 5318585 | http://pubchem.ncbi.nlm.nih.gov/compound/5318585 | PGCKDCPTJAQQSQ-UHFFFAOYSA-N | C20H18O6 | CC(=CCC1=C(C=CC(=C1)C2=C(C(=O)C3=C(C=C(C=C3O2)O)O)O)O)C | CIDm05318585 | CIDs05318585 | isolicoflavonol |
| GANCAO | Glycyrrhiza uralensis | 6801 | http://119.3.41.228:8000/tcmid/herb/6801/ | isoliquiritigenin | 11501 | http://119.3.41.228:8000/tcmid/ingredient/11501/ | 6603886 | http://pubchem.ncbi.nlm.nih.gov/compound/6603886 | DXDRHHKMWQZJHT-BAQGIRSFSA-N | C15H12O4 | C1=CC(=CC=C1C=CC(=O)C2=C(C=C(C=C2)O)O)O | CIDm00000425 | CIDs06603886 | isoliquiritige. |
| GANCAO | Glycyrrhiza uralensis | 6801 | http://119.3.41.228:8000/tcmid/herb/6801/ | isoorientin | 23134 | http://119.3.41.228:8000/tcmid/ingredient/23134/ | 49852298 | http://pubchem.ncbi.nlm.nih.gov/compound/49852298 | ODBRNZZJSYPIDI-VJXVFPJBSA-M | C21H19O11- | C1=CC(=C(C=C1C2=CC(=O)C3=C(C(=C(C=C3O2)O)C4C(C(C(C(O4)CO)O)O)O)[O-])O)O | CIDm01147776 | CIDs00114776 | isoorientin |
| GANCAO | Glycyrrhiza uralensis | 6801 | http://119.3.41.228:8000/tcmid/herb/6801/ | neoisopulegol | 15407 | http://119.3.41.228:8000/tcmid/ingredient/15407/ | 6553885 | http://pubchem.ncbi.nlm.nih.gov/compound/6553885 | ZYTMANIQRDEHIO-UTLUCORTSA-N | C10H18O | CC1CCC(C(C1)O)C(=C)C | CIDm00024585 | CIDs06553885 | isopulegol |
| GANCAO | Glycyrrhiza uralensis | 6801 | http://119.3.41.228:8000/tcmid/herb/6801/ | astragalin | 1935 | http://119.3.41.228:8000/tcmid/ingredient/1935/ | 44258797 | http://pubchem.ncbi.nlm.nih.gov/compound/44258797 | JPUKWEQWGBDDQB-ZVIPYZFUSA-N | C21H20O11 | C1=CC(=CC=C1C2=C(C(=O)C3=C(C=C(C=C3O2)O)O)OC4C(C(C(C(O4)CO)O)O)O)O | CIDm05282102 | CIDs44258797 | kaempferol.-ga. |
| GANCAO | Glycyrrhiza uralensis | 6801 | http://119.3.41.228:8000/tcmid/herb/6801/ | tetrahydroharmine | 21054 | http://119.3.41.228:8000/tcmid/ingredient/21054/ | 442118 | http://pubchem.ncbi.nlm.nih.gov/compound/442118 | ZXLDQJLIBNPEFJ-MRVPVSSYSA-N | C13H16N2O | CC1C2=C(CCN1)C3=C(N2)C=C(C=C3)OC | CIDm00159809 | CIDs00442118 | leptaflorine |
| GANCAO | Glycyrrhiza uralensis | 6801 | http://119.3.41.228:8000/tcmid/herb/6801/ | licoricone | 12788 | http://119.3.41.228:8000/tcmid/ingredient/12788/ | 5319013 | http://pubchem.ncbi.nlm.nih.gov/compound/5319013 | GGWMNTNDTRKETA-UHFFFAOYSA-N | C22H22O6 | CC(=CCC1=C(C=C(C(=C1OC)C2=COC3=C(C2=O)C=CC(=C3)O)O)OC)C | CIDm05319013 | CIDs05319013 | licoricone |
| GANCAO | Glycyrrhiza uralensis | 6801 | http://119.3.41.228:8000/tcmid/herb/6801/ | lensinine | 31440 | http://119.3.41.228:8000/tcmid/ingredient/31440/ | 160644 | http://pubchem.ncbi.nlm.nih.gov/compound/160644 | XCUCMLUTCAKSOZ-FIRIVFDPSA-N | C37H42N2O6 | CN1CCC2=CC(=C(C=C2C1CC3=CC=C(C=C3)O)OC4=C(C=CC(=C4)CC5C6=CC(=C(C=C6CCN5C)OC)OC)O)OC | CIDm01606644 | CIDs00160644 | liensinine |
| GANCAO | Glycyrrhiza uralensis | 6801 | http://119.3.41.228:8000/tcmid/herb/6801/ | liquiritigenin | 12902 | http://119.3.41.228:8000/tcmid/ingredient/12902/ | 928837 | http://pubchem.ncbi.nlm.nih.gov/compound/928837 | FURUXTVZLHCCNA-CQSZACIVSA-N | C15H12O4 | C1C(OC2=C(C1=O)C=CC(=C2)O)C3=CC=C(C=C3)O | CIDm00001889 | CIDs00928837 | liquiritigenin |
| GANCAO | Glycyrrhiza uralensis | 6801 | http://119.3.41.228:8000/tcmid/herb/6801/ | methylglyoxal | 14467 | http://119.3.41.228:8000/tcmid/ingredient/14467/ | 880 | http://pubchem.ncbi.nlm.nih.gov/compound/880 | AIJULSRZWUXGPQ-UHFFFAOYSA-N | C3H4O2 | CC(=O)C=O | CIDm00000880 | CIDs00000880 | methylglyoxal |

| | | | | | | | | | | | | | |
|---|---|---|---|---|---|---|---|---|---|---|---|---|---|
| GANCAO | Glycyrrhiza uralensis | 6801 | http://119.3.41.228:8000/tcmid/herb/6801/ | narwedine | 15280 | http://119.3.41.228:8000/tcmid/ingredient/15280/ | 10446822 | http://pubchem.ncbi.nlm.nih.gov/compound/10446822 | QENVUHCAYXAROT-MCVKWTAESA-N | C17H19NO3 | CN1CCC23C=CC(=O)CC2OC4=C(C=CC(=C34)C1)OC | CIDm00441596 | CIDs10446822 | narwedine |
| GANCAO | Glycyrrhiza uralensis | 6801 | http://119.3.41.228:8000/tcmid/herb/6801/ | tetrahydropalmatine | 21061 | http://119.3.41.228:8000/tcmid/ingredient/21061/ | 6602555 | http://pubchem.ncbi.nlm.nih.gov/compound/6602555 | MGSZZQQRTPWMEI-UHFFFAOYSA-N | C21H26ClNO4 | COC1=C(C2=C(CC3C4=CC(=C(C=C4CCN3C2)OC)OC)C=C1)OC.Cl | CIDm00005417 | CIDs00005417 | tetrahydro.ati. |
| GANCAO | Glycyrrhiza uralensis | 6801 | http://119.3.41.228:8000/tcmid/herb/6801/ | umbelliferone | 22179 | http://119.3.41.228:8000/tcmid/ingredient/22179/ | 45358955 | http://pubchem.ncbi.nlm.nih.gov/compound/45358955 | ORHBXUUXSCNDEV-RALIUCGRSA-N | C9H6O3 | C1=CC(=CC2=C1C=CC(=O)O2)O | CIDm052814426 | CIDs45358955 | umbelliferone |
| HUANGQI | Astragalus membranaceus | 6919 | http://119.3.41.228:8000/tcmid/herb/6919/ | guanosine | 9070 | http://119.3.41.228:8000/tcmid/ingredient/9070/ | 16219429 | http://pubchem.ncbi.nlm.nih.gov/compound/16219429 | YCHNAJLCEKPFHB-GWTDSMLYSA-N | C10H15N5O6 | C1=NC2=C(N1C3C(C(C(O3)CO)O)O)NC(=NC2=O)N.O | CIDm00000765 | CIDs00006802 | 9-beta-D-arabi. |
| HUANGQI | Astragalus membranaceus | 6919 | http://119.3.41.228:8000/tcmid/herb/6919/ | uridine | 22236 | http://119.3.41.228:8000/tcmid/ingredient/22236/ | 45356795 | http://pubchem.ncbi.nlm.nih.gov/compound/45356795 | DRTQHJPVMGBUCF-AYZDMWBASA-N | C9H12N2O6 | C1=CN(C(=O)NC1=O)C2C(C(C(O2)CO)O)O | CIDm00001177 | CIDs45356795 | ara-U |
| HUANGQI | Astragalus membranaceus | 6919 | http://119.3.41.228:8000/tcmid/herb/6919/ | astramembrannin i | 1945 | http://119.3.41.228:8000/tcmid/ingredient/1945/ | 122690 | http://pubchem.ncbi.nlm.nih.gov/compound/122690 | QMNWISYXSJWHRY-XZFBGSIGSA-N | C41H68O14 | CC1(C(CCC23C1C(CC4C2(C3)CCC5(C4(CC(C5C6(CCC(O6)C(C)(C)O)C)O)C)C)OC7C(C(C(C(O7)CO)O)O)O)OC8C(C(C(CO8)O)O)O)C | CIDm00122690 | CIDs00122690 | astragaloside. |
| HUANGQI | Astragalus membranaceus | 6919 | http://119.3.41.228:8000/tcmid/herb/6919/ | glucuronicacid | 8760 | http://119.3.41.228:8000/tcmid/ingredient/8760/ | 441478 | http://pubchem.ncbi.nlm.nih.gov/compound/441478 | AEMOLEFTQBMNLQ-QIUUJYRFSA-N | C6H10O7 | C1(C(C(OC(C1O)O)C(=O)O)O)O)O | CIDm00000610 | CIDs00441478 | beta-D-glu.nid. |
| HUANGQI | Astragalus membranaceus | 6919 | http://119.3.41.228:8000/tcmid/herb/6919/ | β-etaine | 2318 | http://119.3.41.228:8000/tcmid/ingredient/2318/ | 247 | http://pubchem.ncbi.nlm.nih.gov/compound/247 | KWIUHFFTVRNATP-UHFFFAOYSA-N | C5H11NO2 | C[N+](C)(C)CC(=O)[O-] | CIDm00000247 | CIDs00000247 | betaine |
| HUANGQI | Astragalus membranaceus | 6919 | http://119.3.41.228:8000/tcmid/herb/6919/ | calycosin | 3004 | http://119.3.41.228:8000/tcmid/ingredient/3004/ | 5280448 | http://pubchem.ncbi.nlm.nih.gov/compound/5280448 | ZZAJQOPSWWVMBI-UHFFFAOYSA-N | C16H12O5 | COC1=C(C=C(C=C1)C2=COC3=C(C2=O)C=CC(=C3)O)O | CIDm052804448 | CIDs05280448 | calycosin |

| | | | | | | | | | | | | | |
|---|---|---|---|---|---|---|---|---|---|---|---|---|---|
| HUANGQI | Astragalus membranaceus | 6919 | http://119.3.41.228:8000/tcmid/herb/6919/ | canavanine | 3063 | http://119.3.41.228:8000/tcmid/ingredient/3063/ | 25243900 | http://pubchem.ncbi.nlm.nih.gov/compound/25243900 | FSBIGDSBMBYOPN-UHFFFAOYSA-O | C5H13N4O3+ | C(CO[NH+]=C(N)N)C(C(=O)[O-])[NH3+] | CIDm00000275 | CIDs00000275 | canavanine |
| HUANGQI | Astragalus membranaceus | 6919 | http://119.3.41.228:8000/tcmid/herb/6919/ | choline | 3589 | http://119.3.41.228:8000/tcmid/ingredient/3589/ | 305 | http://pubchem.ncbi.nlm.nih.gov/compound/305 | OEYIOHPDSNJKLS-UHFFFAOYSA-N | C5H14NO+ | C[N+](C)(C)CCO | CIDm00000305 | CIDs00000305 | choline |
| HUANGQI | Astragalus membranaceus | 6919 | http://119.3.41.228:8000/tcmid/herb/6919/ | gamma-sitosterol | 29509 | http://119.3.41.228:8000/tcmid/ingredient/29509/ | 636741 | http://pubchem.ncbi.nlm.nih.gov/compound/636741 | KZJWDPNRJALLNS-JQEGLAGPSA-N | C29H50O | CCC(CCC(C)C1CCC2C1(CCC3C2CC=C4C3(CCC(C4)O)C)C)C(C)C | CIDm00006744 | CIDs00636741 | clionasterol |
| HUANGQI | Astragalus membranaceus | 6919 | http://119.3.41.228:8000/tcmid/herb/6919/ | β-sitosterol | 19967 | http://119.3.41.228:8000/tcmid/ingredient/19967/ | 222284 | http://pubchem.ncbi.nlm.nih.gov/compound/222284 | KZJWDPNRJALLNS-VJSFXXLFSA-N | C29H50O | CCC(CCC(C)C1CCC2C1(CCC3C2CC=C4C3(CCC(C4)O)C)C)C(C)C | CIDm00006744 | CIDs00222284 | clionasterol |
| HUANGQI | Astragalus membranaceus | 6919 | http://119.3.41.228:8000/tcmid/herb/6919/ | folic acid | 7851 | http://119.3.41.228:8000/tcmid/ingredient/7851/ | 6037 | http://pubchem.ncbi.nlm.nih.gov/compound/6037 | OVBPIULPVIDEAO-LBPRGKRZSA-N | C19H19N7O6 | C1=CC(=CC=C1C(=O)NC(CCC(=O)O)C(=O)O)NCC2=CN=C3C(=N2)C(=O)N=C(N3)N | CIDm00003405 | CIDs00006037 | folic acid |
| HUANGQI | Astragalus membranaceus | 6919 | http://119.3.41.228:8000/tcmid/herb/6919/ | formononetin | 7882 | http://119.3.41.228:8000/tcmid/ingredient/7882/ | 10378473 | http://pubchem.ncbi.nlm.nih.gov/compound/10378473 | HKQYGTCOTHHOMP-OPOMHRLSSA-N | C16H12O4 | COC1=CC=C(C=C1)C2=COC3=C(C2=O)C=CC(=C3)O | CIDm05280378 | CIDs10378473 | formononetin |
| HUANGQI | Astragalus membranaceus | 6919 | http://119.3.41.228:8000/tcmid/herb/6919/ | isorhamnetin | 11645 | http://119.3.41.228:8000/tcmid/ingredient/11645/ | 25202413 | http://pubchem.ncbi.nlm.nih.gov/compound/25202413 | IZQSVPBOUDKVDZ-UHFFFAOYSA-M | C16H11O7- | COC1=C(C=CC(=C1)C2=C(C(=O)C3=C(C=C(C=C3O2)O)O)[O-])O | CIDm05281654 | CIDs05281654 | isorhamnetin |
| HUANGQI | Astragalus membranaceus | 6919 | http://119.3.41.228:8000/tcmid/herb/6919/ | kumatakenin | 12327 | http://119.3.41.228:8000/tcmid/ingredient/12327/ | 5318869 | http://pubchem.ncbi.nlm.nih.gov/compound/5318869 | BJBUTJQYZDYRMJ-UHFFFAOYSA-N | C17H14O6 | COC1=CC(=C2C(=C1)OC(=C(C2=O)OC)C3=CC=C(C=C3)O)O | CIDm05318869 | CIDs05318869 | kumatakenin |
| HUANGQI | Astragalus membranac | 6919 | http://119.3.41.228:8000/tcmid/herb/6919/ | folinic acid | 7852 | http://119.3.41.228:8000/tcmid/ingredient/7852/ | 149436 | http://pubchem.ncbi.nlm.nih.gov/compound/149436 | VVIAGPKUTFNRDU-STQMWFEESA-N | C20H23N7O7 | C1C(N(C2=C(N1)NC(=NC2=O)N)C=O)CNC3=CC=C(C=C3)C(=O)NC(CCC(=O)O)C(=O)O | CIDm00000143 | CIDs00149436 | leucovorin |

| | | | | | | | | | | | | | | |
|---|---|---|---|---|---|---|---|---|---|---|---|---|---|---|
| | eus | | | | | | | | | | | | | |
| HUANGQI | Astragalus membranaceus | 6919 | http://119.3.41.228:8000/tcmid/herb/6919/ | (+)-medicarpin | 13631 | http://119.3.41.228:8000/tcmid/ingredient/13631/ | 73067 | http://pubchem.ncbi.nlm.nih.gov/compound/73067 | NSRJSISNDPOJOP-CZUORRHYSA-N | $C_{16}H_{14}O_4$ | COC1=CC2=C(C=C1)C3COC4=C(C3O2)C=CC(=C4)O | CIDm00073067 | CIDs00073067 | medicarpin |
| HUANGQI | Astragalus membranaceus | 6919 | http://119.3.41.228:8000/tcmid/herb/6919/ | quercetin | 18302 | http://119.3.41.228:8000/tcmid/ingredient/18302/ | 46906036 | http://pubchem.ncbi.nlm.nih.gov/compound/46906036 | REFJWTPEDVJJIY-UHFFFAOYSA-M | $C_{15}H_9O_7^-$ | C1=CC(=C(C=C1C2=C(C(=O)C3=C(C=C(C=C3O2)O)O)[O-])O)O | CIDm05280343 | CIDs05280343 | quercetin |
| HUANGQI | Astragalus membranaceus | 6919 | http://119.3.41.228:8000/tcmid/herb/6919/ | soyasaponin 1 | 24621 | http://119.3.41.228:8000/tcmid/ingredient/24621/ | 122097 | http://pubchem.ncbi.nlm.nih.gov/compound/122097 | PTDAHAWQAGSZDD-IOVCITQVSA-N | $C_{48}H_{78}O_{18}$ | CC1C(C(C(C(O1)OC2C(C(C(OC2OC3C(C(C(OC3OC4CCC5(C(C4(C)CO)CCC6(C5CC=C7C6(CCC8(C7CC(CC8O)(C)C)C)C)C)C)C(=O)O)O)O)CO)O)O)O)O)O | CIDm00122097 | CIDs00122097 | soyasaponin I |
| HUANGQI | Astragalus membranaceus | 6919 | http://119.3.41.228:8000/tcmid/herb/6919/ | sucrose | 20430 | http://119.3.41.228:8000/tcmid/ingredient/20430/ | 53301851 | http://pubchem.ncbi.nlm.nih.gov/compound/53301851 | CZMRCDWAGMRECN-FXFGBNHJSA-N | $C_{12}H_{22}O_{11}$ | C(C1C(C(C(C(O1)OC2(C(C(C(O2)CO)O)O)CO)O)O)O)O | CIDm00001115 | CIDs53301851 | sucrose |
| HUANGQI | Astragalus membranaceus | 6919 | http://119.3.41.228:8000/tcmid/herb/6919/ | lupeol | 13091 | http://119.3.41.228:8000/tcmid/ingredient/13091/ | 44586882 | http://pubchem.ncbi.nlm.nih.gov/compound/44586882 | MQYXUWHLBZFQQO-CZYRXZFSSA-N | $C_{30}H_{50}O$ | CC(=C)C1CCC2(C1C3CCC4C5(CCC(C(C5CCC4(C3(CC2)C)C)(C)C)O)C)C | CIDm00073040 | CIDs44586882 | triterpenelup. |
| HUANGQI | Astragalus membranaceus | 6919 | http://119.3.41.228:8000/tcmid/herb/6919/ | adenine nucleoside | 618 | http://119.3.41.228:8000/tcmid/ingredient/618/ | 60961 | http://pubchem.ncbi.nlm.nih.gov/compound/60961 | OIRDTQYFTABQOQ-KQYNXXCUSA-N | $C_{10}H_{13}N_5O_4$ | C1=NC2=C(C(=N1)N)N=CN2C3C(C(C(O3)CO)O)O | CIDm00000191 | CIDs00060961 | vidarabine |
